# Vaccination decreases the risk of influenza A virus reassortment with a concomitant increase in subgenomic genetic variation in pigs

**DOI:** 10.1101/2022.03.31.486502

**Authors:** Chong Li, Marie R. Culhane, Declan C. Schroeder, Maxim C-J. Cheeran, Lucina Galina Pantoja, Micah L. Jansen, Montserrat Torremorell

## Abstract

Although vaccination is broadly used in North American swine breeding herds, managing swine influenza is challenging primarily due to the continuous evolution of influenza A virus (IAV) and the ability of the virus to transmit among vaccinated pigs. Studies that have simultaneously assessed the impact of vaccination on the emergence of IAV reassortment and genetic variation in pigs are limited. Here we directly sequenced 28 bronchoalveolar lavage fluid (BALF) samples collected from vaccinated and unvaccinated pigs co-infected with H1N1 and H3N2 IAV strains, and characterized 202 individual viral plaques recovered from 13 BALF samples. We identified 54 reassortant viruses that were grouped in 16 distinct and 18 mixed genotypes. Notably, we found that prime-boost vaccinated pigs had less reassortant viruses than non-vaccinated pigs, likely due to a reduction in the number of days pigs were co-infected with both challenge viruses. However, direct sequencing from BALF samples revealed limited impact of vaccination on viral variant frequency, evolutionary rates, and nucleotide diversity in any IAV coding regions. Overall, our results highlight the value of IAV vaccination not only at limiting virus replication in pigs but also at protecting public health by restricting the generation of novel reassortants with zoonotic and/or pandemic potential.

## Introduction

Influenza A viruses (IAV) are important respiratory pathogens in both humans and pigs globally. In the United States (US), IAV infection causes a significant disease burden on the healthcare system and society, resuling in 12,000 to 61,000 human deaths plus estimated losses of $11.2 billion annually [1,2]. IAV infections in pigs are also considered one of the top disease concerns for the US swine industry when influenza-induced respiratory disease severely reduces pig health and subsequently the pork producers’ profitability and sustainability. Human and swine share the same IAV subtypes, which have similar tangled evolutionary histories contributing to the bidirectional transmission of IAV between both species [3]. Pigs pose a risk for generating novel IAV strains with zoonotic and pandemic potential, which represents an unpredictable threat to both the swine industry and global public health.

IAV exhibits an extraordinary ability for cross-species transmission and immune evasion by continually expanding its genetic diversity through antigenic drift and shift. This genetic diversity permits the rapid evolution of IAV, maximizing the virus’s opportunity to remain viable following significant changes in the environment and making the use of vaccines extremely difficult for disease prevention [4–6]. Therefore, minimizing IAV diversity should be a key strategy for One Health purposes to efficiently control IAV transmission between humans and pigs [7]. As swine are susceptible to avian, human, and swine-origin influenza virus, IAV introduction from multiple hosts immensely enriches the genetic pool of swine IAV and is responsible for the emergence of distinct H1 and H3 IAV lineages during the last 20 years in pigs [8,9]. The distribution of IAV receptors in the swine respiratory tract also promotes IAV co-infections with strains from various hosts and facilitates virus reassortment that may result in new viruses [10]. A mathematical model showed that about 16.8% of IAV co-infection events will lead to virus reassortment in French pigs [11]. This estimate seems reasonable since the recurrent and co-circulation of distinct IAV subtypes within swine herds is quite common and over 74 different H1 genotypes have been detected in the US pig population alone from 2009 to 2016 [12–14]. With such diverse viral populations, swine should be considered one of the potential sources for novel IAV variants of zoonotic and pandemic disease. The 2009 H1N1 virus (pdm09) which is a reassortant virus that originated in pigs, contained gene segments of avian, swine, and human influenza viruses which led to the first influenza pandemic of the 21st century [15]. Within the first year of circulation, between 151,700 and 575,400 people died worldwide due to the 2009 H1N1 virus infection [16]. Due to the continued evolution, the pdm09 virus has generated a complex hemaggulminin (HA) clade system (https://nextstrain.org/flu/seasonal/h1n1pdm/ha/12y) [17]. Moreover, the pdm09 virus has further reassorted with endemic IAV strains in pigs and has generated reassortants with distinct genetic constellations in many countries [18–22]. Some reassortants of specific genotypes have already become the predominant circulating strains in swine populations and have caused fatal infections in people in contact with pigs [23,24].

Even though vaccination is widely used in US pig farms to limit the impact of influenza infections, IAV continues to be a problem in pigs since it evolves rapidly to escape host immunity and transmits under immune conditions [25]. Therefore, understanding how immunity induced by swine IAV vaccination shapes within-host virus evolution in pigs is key to controlling the disease and the emergence of novel antigenic variants. Previous studies have characterized the IAV mutational spectra within naïve vs. vaccinated pigs and other mammals using experimental transmission models which simulated the impact of immune pressure on within-host diversity [25–27]. However, *in vivo* studies that explore how vaccination impacts reassortment between multiple subtypes of IAVs in pigs are lacking. Besides, most of the knowledge on IAV within-host diversity in pigs so far is based on samples taken from nasal cavities and studies that quantified the IAV within-host variation in pig lungs are lacking [25,28]. Considering that the tissue tropism plays a vital role in IAV dissemination along the swine respiratory tract [29], with the pig lungs harboring IAV populations with the most extensive genomic variations, the effect of immune pressure on the IAV within-host diversity may be differ by anatomical location [29–31]. Therefore, *in vivo* studies are needed to provide an integrated picture of how vaccine-induced immunity affects IAV evolutionary trajectories occurring in the swine lower respiratory tract by evaluating the extent of IAV reassortment and mutational spectra taking place concurrently in naïve and vaccinated pigs.

We previously published a vaccine-challenge study assessing IAV infections in pigs vaccinated with five distinct prime-boost vaccine combinations after simultaneous infection with both an H1N1 and an H3N2 IAV strain using a seeder pig model [32]. The bronchoalveolar lavage fluid (BALF) samples obtained from the aforementioned study enabled us to evaluate, in the present study, how IAV reassorts and mutates in the swine lower respiratory tract under immune pressure. We hypothesized that the dual-subtype IAV co-infection model better represents the conditions of IAV co-infection encountered in the field and that the findings could contribute to the body of knowledge of within-host virus evolution in pigs [33]. Here, we performed next-generation sequencing directly on BALF samples and IAV plaques purified from the BALF samples to identify the virus mutations and reassortment that happened in swine lungs.

## Results

### Influenza co-infection model and specimen collection

The BALF specimens utilized in this study originated from a previoulsy published vaccine-challenge study [32]. We used up a co-infection challenge model by inoculating 14 naïve pigs, i.e., seeder pigs, with either an H1N1 (A/swine/Minnesota/PAH-618/2011) or an H3N2 (A/swine/Minnesota/080470/2015) IAV strain and distributed these seeder pigs into 2 rooms by strain (e.g.., seven H1N1 inoculated pigs were housed together in one room and seven H3N2 inoculated were housed together in another room, etc.). After all the seeder pigs were confirmed shedding IAV at two days post-inoculation, two seeder pigs, one inoculated with an H1N1 and the other one with an H3N2 were commingled with ten other pigs in each new room to mimic simultaneous exposure to both strains and subsequent co-infection. As a result, the 14 seeder pigs were distributed as pairs into 7 rooms. In-contact pigs had been vaccinated using different vaccine combinations that included a commercial (COM) multivalent whole inactivated vaccine (WIV), an autogenous (AUT) multivalent WIV, or a bivalent live attenuated influenza vaccine (LAIV). The seeder pigs served as the infection source to the treatment pigs (Figure 1). Fifty treatment pigs that received five different inactivated vaccine combinations (including COM/COM, AUT/AUT, AUT/COM, COM/AUT, and NO VAC/CHALL) were evenly distributed into 5 rooms (two pigs/ treatment/ room). Another twenty pigs that received two different administrations of the LAIV, including LAIV/COM and LAIV/NONE were distributed evenly into two other rooms (five pigs/ treatment/ room). The BALF samples from all the treatment pigs were collected at necropsy seven days post contact with seeder pigs. We obtained IAV genome data from 28 BALF samples from the direct sequencing to identify IAV mutations (Table 1).

**Figure 1.**
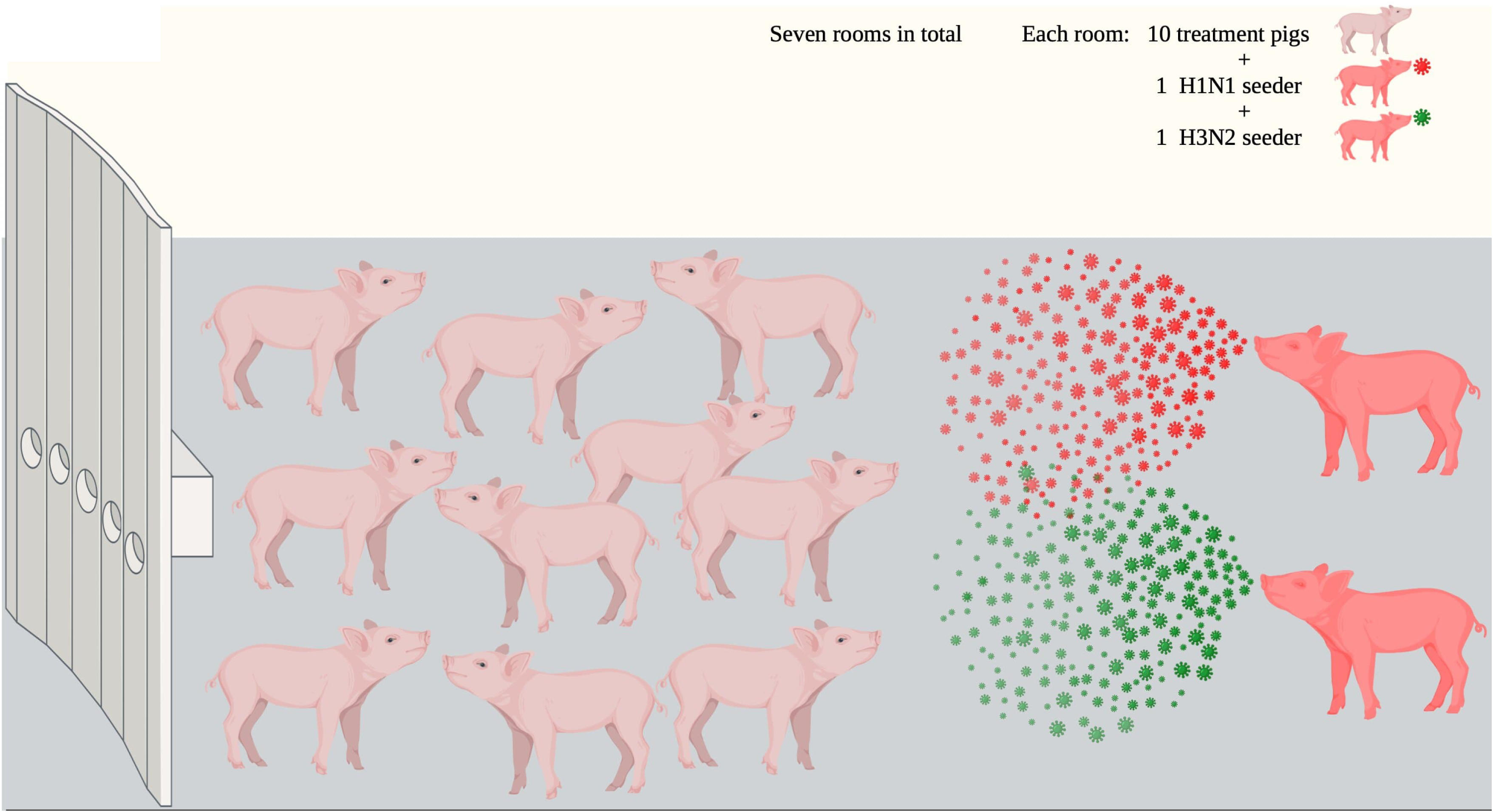
Diagram showing the seeder pig infection model. Fourteen naïve pigs (seeder pigs) were either challenged using an H1N1 or an H3N2 virus and evenly distributed in pairs into seven rooms at approximately 48 hours post-inoculation. From each room, there are two seeder pigs were commingled with ten treatment pigs that had been vaccinated using different vaccine combinations of a commercial multivalent whole inactivated vaccine (COM), an autogenous multivalent whole inactivated vaccine (AUT), or a bivalent live attenuated vaccine (LAIV), and served as the infection source to treatment pigs. BALF samples from all treatment pigs were collected at necropsy seven days post contact with the seeder pigs. Treatment pigs were grouped into PRIME BOOST, SINGLE LAIV, and NO VAC based on the vaccine doses they received. The diagram was adapted from Li et al., 2020 and created by BioRender.com.

**Table 1.**
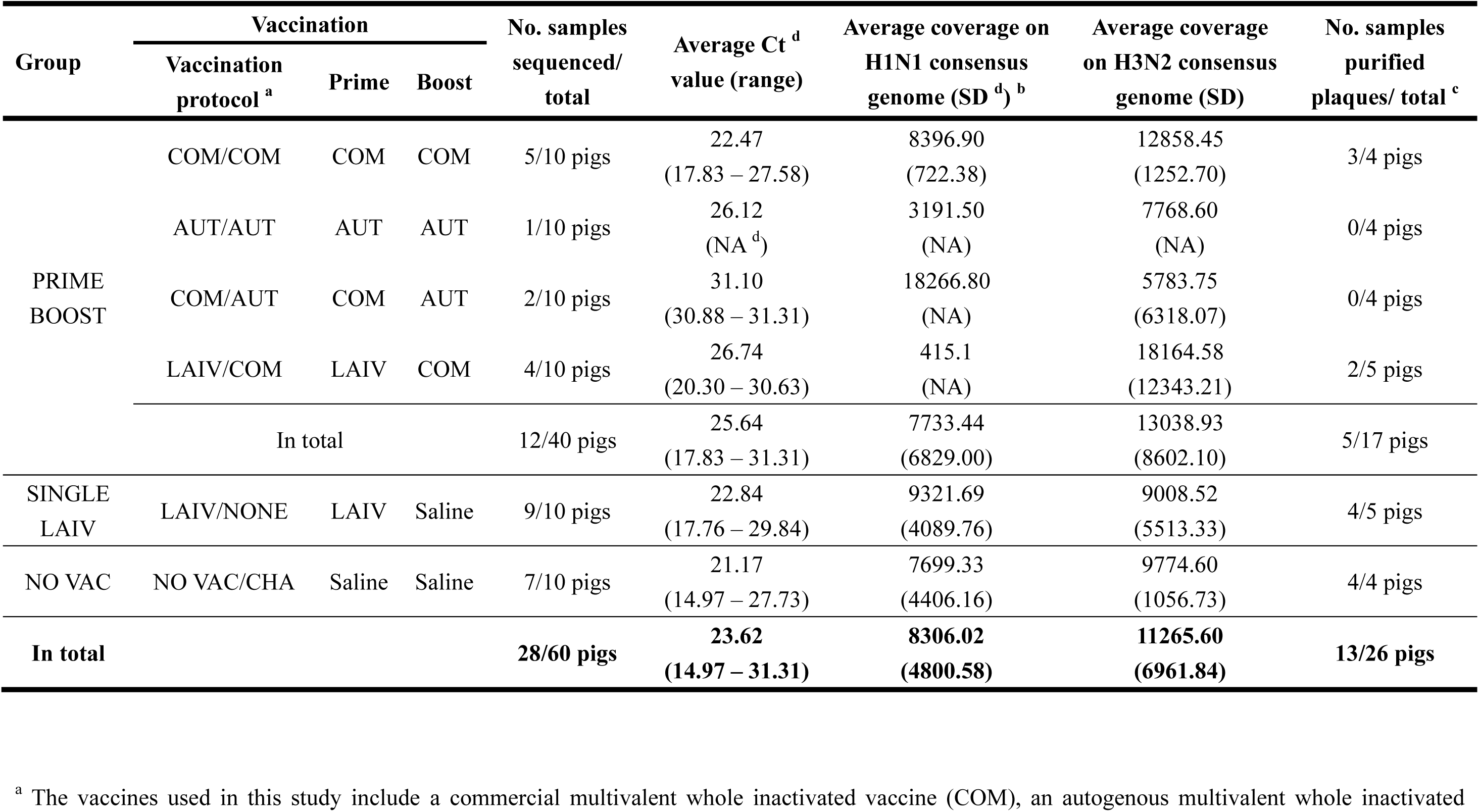
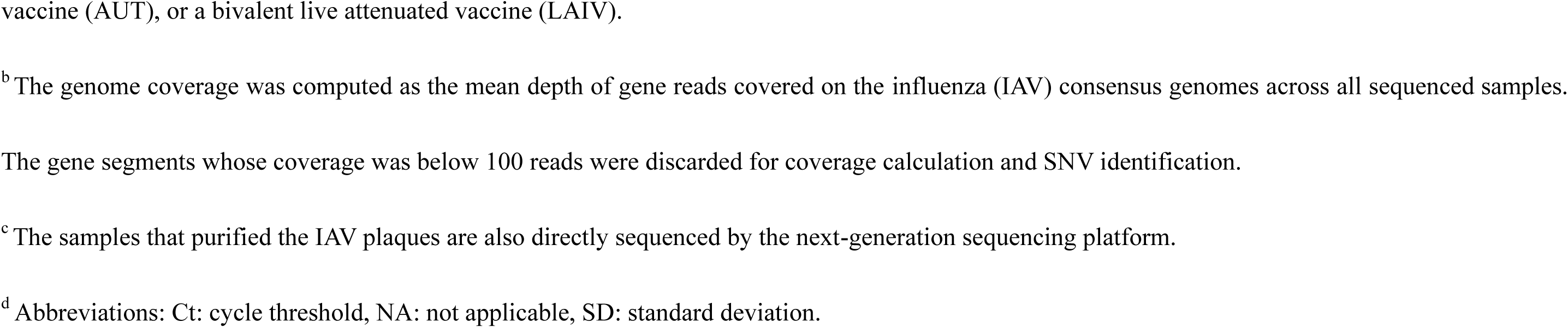
Number of bronchoalveolar lavage fluid (BALF) samples available for direct sequencing or plaque purification from each group.

| Group | Vaccination |  |  | No. samples sequenced/ total | Average Ct <sup>d</sup> value (range) | Average coverage on H1N1 consensus genome (SD <sup>d</sup>) <sup>b</sup> | Average coverage on H3N2 consensus genome (SD) | No. samples purified plaques/ total <sup>c</sup> |
| --- | --- | --- | --- | --- | --- | --- | --- | --- |
|  | Vaccination protocol <sup>a</sup> | Prime | Boost |  |  |  |  |  |
| PRIME BOOST | COM/COM | COM | COM | 5/10 pigs | 22.47<br>(17.83 – 27.58) | 8396.90<br>(722.38) | 12858.45<br>(1252.70) | 3/4 pigs |
|  | AUT/AUT | AUT | AUT | 1/10 pigs | 26.12<br>(NA <sup>d</sup>) | 3191.50<br>(NA) | 7768.60<br>(NA) | 0/4 pigs |
|  | COM/AUT | COM | AUT | 2/10 pigs | 31.10<br>(30.88 – 31.31) | 18266.80<br>(NA) | 5783.75<br>(6318.07) | 0/4 pigs |
|  | LAIV/COM | LAIV | COM | 4/10 pigs | 26.74<br>(20.30 – 30.63) | 415.1<br>(NA) | 18164.58<br>(12343.21) | 2/5 pigs |
|  | In total |  |  | 12/40 pigs | 25.64<br>(17.83 – 31.31) | 7733.44<br>(6829.00) | 13038.93<br>(8602.10) | 5/17 pigs |
| SINGLE LAIV | LAIV/NONE | LAIV | Saline | 9/10 pigs | 22.84<br>(17.76 – 29.84) | 9321.69<br>(4089.76) | 9008.52<br>(5513.33) | 4/5 pigs |
| NO VAC | NO VAC/CHA | Saline | Saline | 7/10 pigs | 21.17<br>(14.97 – 27.73) | 7699.33<br>(4406.16) | 9774.60<br>(1056.73) | 4/4 pigs |
| **In total** |  |  |  | **28/60 pigs** | **23.62**<br>**(14.97 – 31.31)** | **8306.02**<br>**(4800.58)** | **11265.60**<br>**(6961.84)** | **13/26 pigs** |

<sup>a</sup> The vaccines used in this study include a commercial multivalent whole inactivated vaccine (COM), an autogenous multivalent whole inactivated
vaccine (AUT), or a bivalent live attenuated vaccine (LAIV).
<sup>b</sup> The genome coverage was computed as the mean depth of gene reads covered on the influenza (IAV) consensus genomes across all sequenced samples.
The gene segments whose coverage was below 100 reads were discarded for coverage calculation and SNV identification.
<sup>c</sup> The samples that purified the IAV plaques are also directly sequenced by the next-generation sequencing platform.
<sup>d</sup> Abbreviations: Ct: cycle threshold, NA: not applicable, SD: standard deviation.

To explore the IAV reassortment in vaccinated and non-vaccinated pigs, we selected pigs in three rooms (two rooms containing WIV treatment pigs and one room containing LAIV treatment pigs) within which seeder and treatment pigs were confirmed to be shedding virus in nasal secretions rRT-PCR. Both, H1N1 and H3N2 seeders in the selected rooms shed IAV for at least two days after being mixed with the treatment pigs. All the BALF samples from treatment pigs confirmed virus isolation positive were further investigated using the plaque assay. Finally, 13 BALF samples yielded IAV plaques (Table 1). Based on the vaccination regime, treatment pigs were grouped into PRIME BOOST (with COM/COM, AUT/AUT, COM/AUT, and LAIV/COM pigs), SINGLE LAIV (with LAIV/NONE pigs), and NO VAC (with NO VAC/CHALL pigs) groups and used for further analysis. There were no samples from AUT/COM vaccinated pigs, since that WIV vaccine treatment prevented shedding and no virus was isolated from any pig in that treatment group.

### Multiple genotypes identified among the new reassortant, plaque-purified viruses

To characterize and evaluate the distribution of reassortant viruses in vaccinated and non-vaccinated pigs, a total of 202 IAV plaques were isolated and whole-genome sequenced from 13 BALF samples collected at necropsy from the pigs receiving the prime-boost (PRIME BOOST), single-dose LAIV (SINGLE LAIV), and no vaccine (NO VAC) administrations (Supplementary file 1). A summary of the genotypes is shown in Figure 1. Among the 202 plaques, 148 (73.27%) were the parental virus challenge strains (137 (67.82%) H3N2 and 11 (5.45%) H1N1). Fifty-four (27%) plaques were classified as IAV reassortants with 34 (16.83%) plaques distributed into 16 distinct gene segment constellations (R01-R16) or genotypes, and 20 (9.90%) plaques classified as 18 mixed genotypes (M01-M18) (Figure 2 and Figure 2 – source data 1). Mixed genotypes contained complete gene sequences of both parental viruses in a given gene segment. The IAV reassortants were detected in six out of 13 pigs with some pigs having as few as one genotype and as many as 14 genotypes (including mixed genotypes); notably 85% (46/54) of reassortants originated from only three pigs (Figure 2 – figure supplement 1).

**Figure 2.**
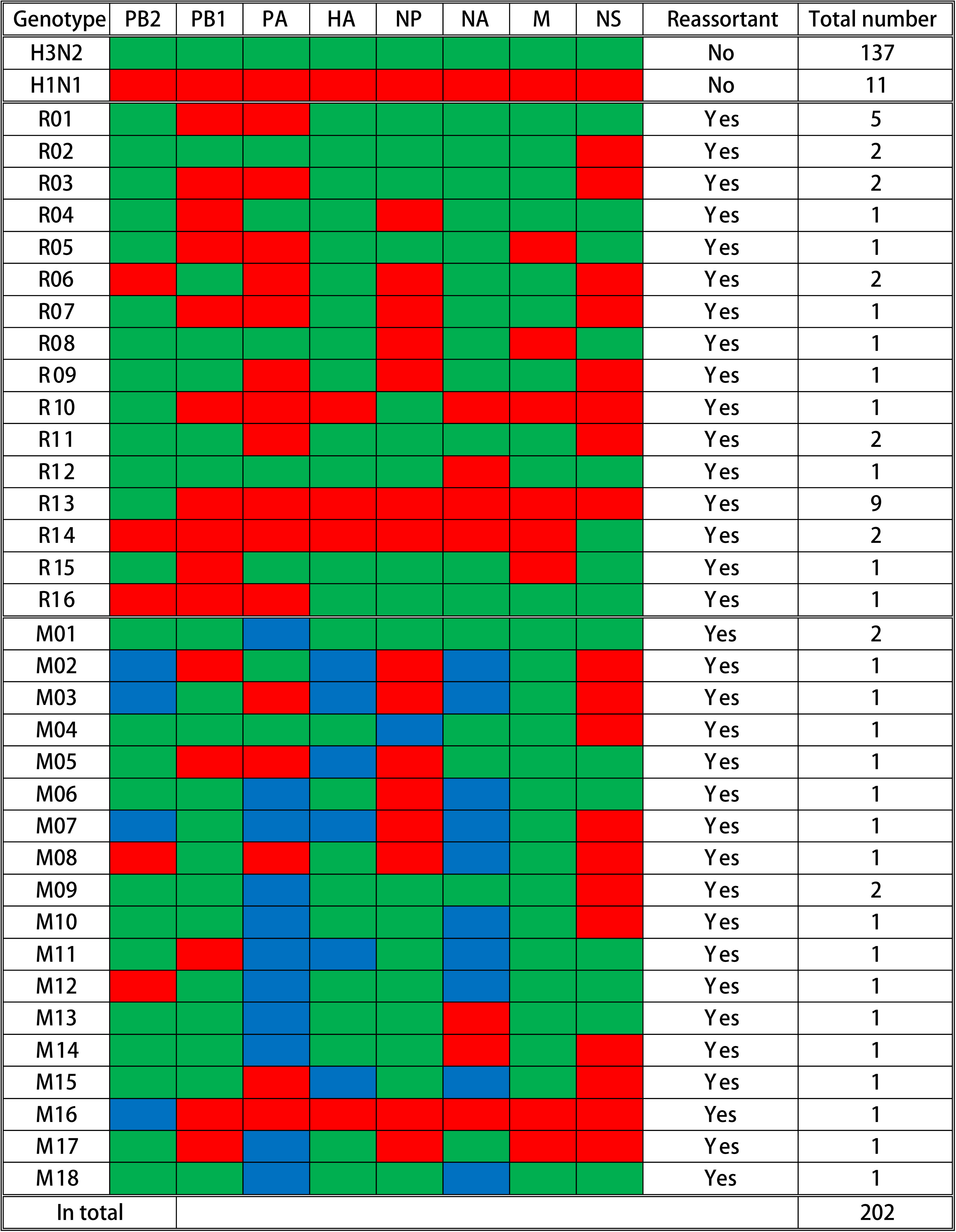
Summary of genotypes detected in the influenza A virus plaques. A total of 202 plaques were whole genome sequenced and genotyped based on the Illumina Nextseq platform. Gene segments are shown above the columns. Red blocks represent gene segments that originate from the H1N1 virus; green blocks originate from the H3N2 virus; and blue blocks indicate that complete gene segments were detected from both viruses. The specific genotype number is indicated on the left side of each row, and the quantity of plaques that contain the corresponding gene constellation is shown on the right side of the row. The specific reassortant genotype number is named after R, and the M-number indicates the specific mixed genotype number. The quantity and genotypes of influenza plaques isolated from each individual pig are displayed in Figure 2 – figure supplement 1. The maximum likelihood trees and the assembled nucleotide sequences of isolated plaques used for constructing the trees can be found in Figure 2 – source data 1 and Figure 2 – source data 2, respectively.

### Vaccination decreases the number of reassortant influenza A viruses

To evaluate whether vaccination alters the occurrence of IAV reassortment in swine lungs, we compared the percentage of IAV reassortants isolated from vaccinated and non-vaccinated pigs (Figure 3A). Non-vaccinated (NO VAC) pigs had more reassortant viruses and more distinct genotypes, with 54% (37/74) of plaques being reassortants which belong to 10 single and 17 mixed genotypes. For the remaining plaques isolated in PRIME BOOST and SINGLE LAIV pigs, 13.2% (17/128) were reassortants distributed in eight single and three mixed genotypes. We found 25.0% (13/52) of plaques isolated from the SINGLE LAIV pigs and 5.41% (4/74) of plaques in the PRIME BOOST pigs were identified as reassortants. The proportion of reassortants was significantly lower in the PRIME BOOST pigs than the NO VAC pigs (p=0.022).

**Figure 3.**
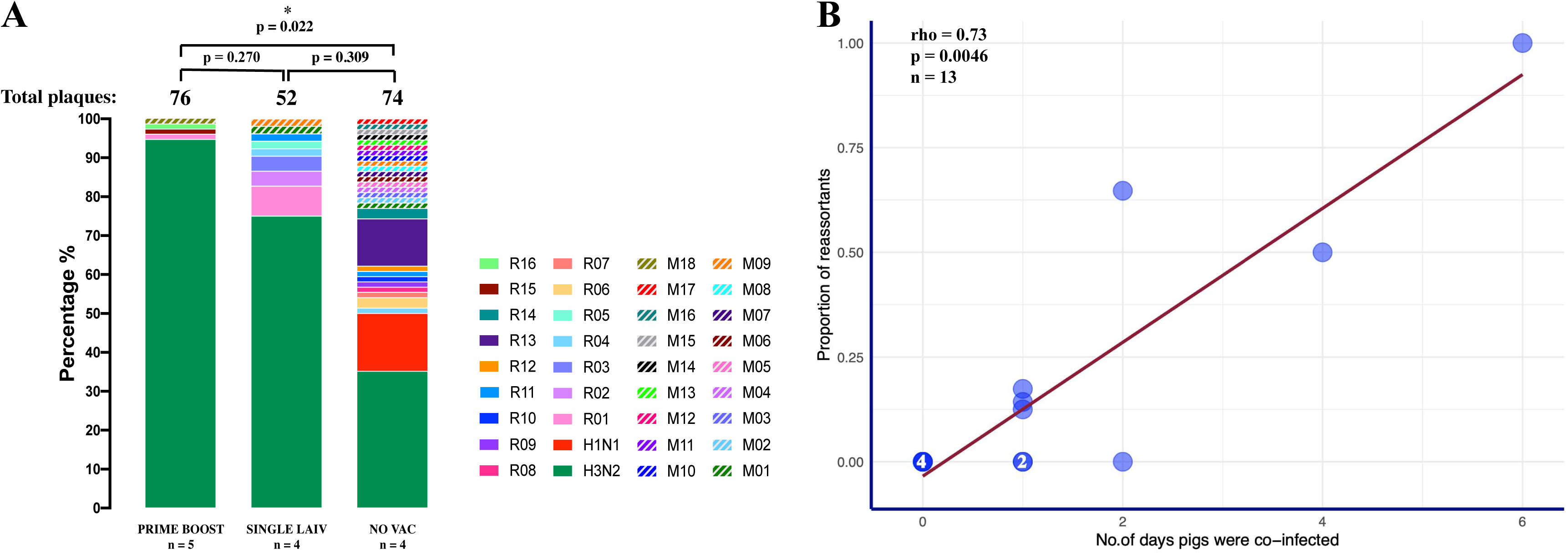
Emergence of reassortant influenza A viruses (IAV) is correlated with the number of days that pigs are co-infected with H1 and H3 viruses. (A) Percentage of reassortant plaques in pigs by genotype and treatment groups. Each genotype is shown in a different color. The total number of plaques for each group is shown above each bar, and the quantity of available BALF (broncho alveolar lavage fluid) samples for each group (n) is indicated under the treatment names. The proportion of IAV reassortants was compared by the binomial logistic regression model, allowing for overdispersion. P-value < 0.05 was considered significant. (B) Correlation between the proportion of reassortant viruses and the number of days pigs were co-infected with H1 and H3 challenge viruses. The number in the dot represents the number of overlapping points plotted for pigs that had the same proportion of reassortants and co-infection day, and the total number of samples available for this analysis is indicated (n). The number of days pigs were co-infected is shown in Figure 3 – source data 1, which defined as the number of days when both H1 and H3 IAV were detected in the nasal cavities or lungs by a HA subtype specific multiplex rRT-PCR. Spearman’s Rank-Order Correlation test evaluated the direction and intensity of the correlation between the proportion of reassortant viruses and the number of days pigs were co-infected.

To further investigate whether the occurrence of reassortant viruses increased with increasing length of time (in days) that pigs were co-infected, we evaluated the relationship between the duration of co-infection (i.e., number of days the pigs were co-infected with H1N1 and H3N2 viruses), as measured by a subtype specific multiplex rRT-PCR in nasal swabs and BALF samples collected at 2 to 7 dpc and the total number of reassortants detected in BALF at necropsy performed on 7dpc (Figure 3 – source data 1). We observed a strong positive correlation (R=0.73, p=0.0046) as evaluated by the Spearman’s Rank-Order Correlation test, between the proportion of reassortant viruses isolated with an increasing duration of co-infection (Figure 3B).

### Virus load and genome coverage of direct sequenced BALF samples for single nucleotide variant (SNV) identification

Virus load and sequencing coverage are considered important factors that could affect the accuracy of IAV SNV identification [34]. The sequenced samples in this study had a high virus load with a mean Ct value of 23.62 (range 14.97 to 31.31) tested by IAV matrix real-time PCR (rRT-PCR). The average Ct values of sequenced samples were not significantly different between treatment groups (p = 0.13, Kruskal-Wallis rank-sum test). We excluded the sequences from any IAV segments whose average coverages were less than 100 reads [35]. The mean depth across the whole available samples used for SNV identification in the H1N1 and H3N2 genomes are 8306.02 and 11265.60 reads, respectively (Table 1). The mean genome coverages on H1N1 and H3N2 viruses from the sequenced samples were not significantly different between treatment groups (H1N1: p = 0.49, H3N2: p = 0.46, Kruskal-Wallis rank-sum test).

### Vaccine-induced immunity has limited impact on within-pig HA nucleotide variation of H1N1 and H3N2 viruses

We identified within-host viral variants from BALF samples separately in H1N1 (A/swine/Minnesota/PAH-618/2011) and H3N2 (A/swine/Minnesota/080470/2015) challenged pigs based on the single nucleotide variants (SNVs) selection criteria described in Methods. The SNVs identified in H1N1 and H3N2 coding regions were reported based on the H1 and H3 numbering schemes, respectively, included the signal peptide.

We called a total of 380 SNVs in 219,659 sequenced H1N1 consensus nucleotides (206 nonsynonymous, 166 synonymous, 8 stop-gained) from 19 BALF samples (Figure 4 - figure supplement 1A and Figure 4 – source data 1). H1N1 SNVs were dominated by low-frequency variants (Figure 3A). About 9.5% (36/380) of SNVs existed at the consensus level (above 50% of virus population) and over 81.8% (311/380) of SNVs, which included 170 (82.5%) nonsynonymous, 133 (80.1%) synonymous, and 8 (100.0%) stop-gained SNVs, were present at less than 10% frequency. The average number and frequency of H1N1 SNVs by gene segment and group are summarized in Table 2. The quantity and frequency of SNVs in each IAV gene did not differ significantly among treatment groups (p = 0.06 – 0.99 for SNVs quantity; p = 0.06 – 0.80 for SNVs frequency, Kruskal-Wallis rank-sum test). The H1N1 virus exhibited low antigenic variation regardless of treatment groups (Figure 4B). Only two synonymous nucleotide polymorphisms were found in the H1 Sb antigenic region (H1 nucleotide site 636) within the same pig from the PRIME BOOST group at extremely low frequency (< 4%). The number of H1 SNVs did not differ based on the intensity of the humoral and cellular immunity induced by vaccination (Figures 4C and 4D). There was no evidence of correlation of the H1 nucleotide divergence with the H1 specific HI titers (R=-0.069, p=0.79, Spearman’s Rank-Order Correlation test) nor with the H1 specific IFN-γ secreting cell spots (R=-0.153, p=0.62, Spearman’s Rank-Order Correlation test).

**Figure 4.**
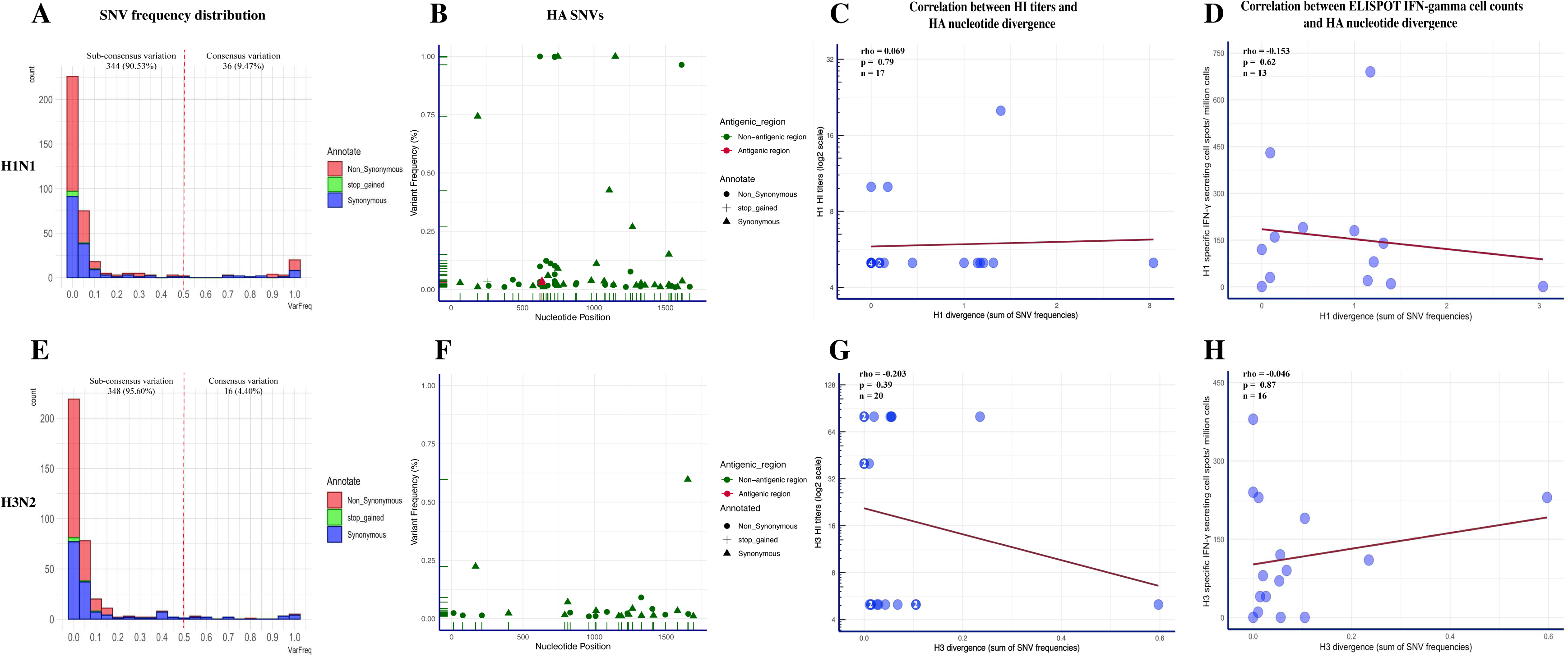
Summary of single nucleotide variants (SNV) frequency and HA nucleotide variations of H1N1 and H3N2 virus within pigs from different vaccination statuses. The frequency distribution of the H1N1 (A) and H3N2 (E) SNVs in pigs regardless of treatment groups. The quantity of SNVs at a given frequency interval (bin width = 0.05) is presented by a stacked histogram based on the mutation types. The SNVs with frequencies above 0.5 were considered as consensus variants. Antigenic variant identification from the total detected H1(B) and H3 (F) SNVs. The HA SNVs are colored based on whether they fell into the H1 antigenic regions (include the Sa, Sb, Ca1, Ca2 and Cb region) or H3 antigenic regions (include the A, B, C, D and E region). Only two H1 SNVs were identified in the Sb region (red) and the rest of the H1 SNVs were in non-antigenic regions (green). For H3 SNVs, all the detected SNVs were in non-antigenic regions (green). The Spearman correlation between H1 specific HI titer (C) or H1 specific IFN-γsecreting cell counts (D) with the H1 nucleotide divergence identified in individual pigs were calculated. The same statistics were also computed for H3 viruses between the H3 specific HI titer (G) or H3 specific IFN-γsecreting cell counts (H) with the H3 nucleotide divergence identified in individual pigs. HA nucleotide divergence was calculated by summing the frequencies of all the HA SNVs identified in each sample. Same as Fig 3B, the number in specific dots represents the quantity of overlapping points plotted for pigs that had the same numbers for both variables, and the total quantity of available samples for this analysis is indicated as “n”. All the within-host IAV nucleotide polymorphisms present in at least 1% of H1N1 and H3N2 sequencing reads from BALF samples are shown in Figure 4 - figure supplement 1. The detailed information of identified H1N1 and H3N2 SNVs are displayed in Figure 4 – source data 1 and Figure 4 – source data 2, respectively.

**Table 2.**
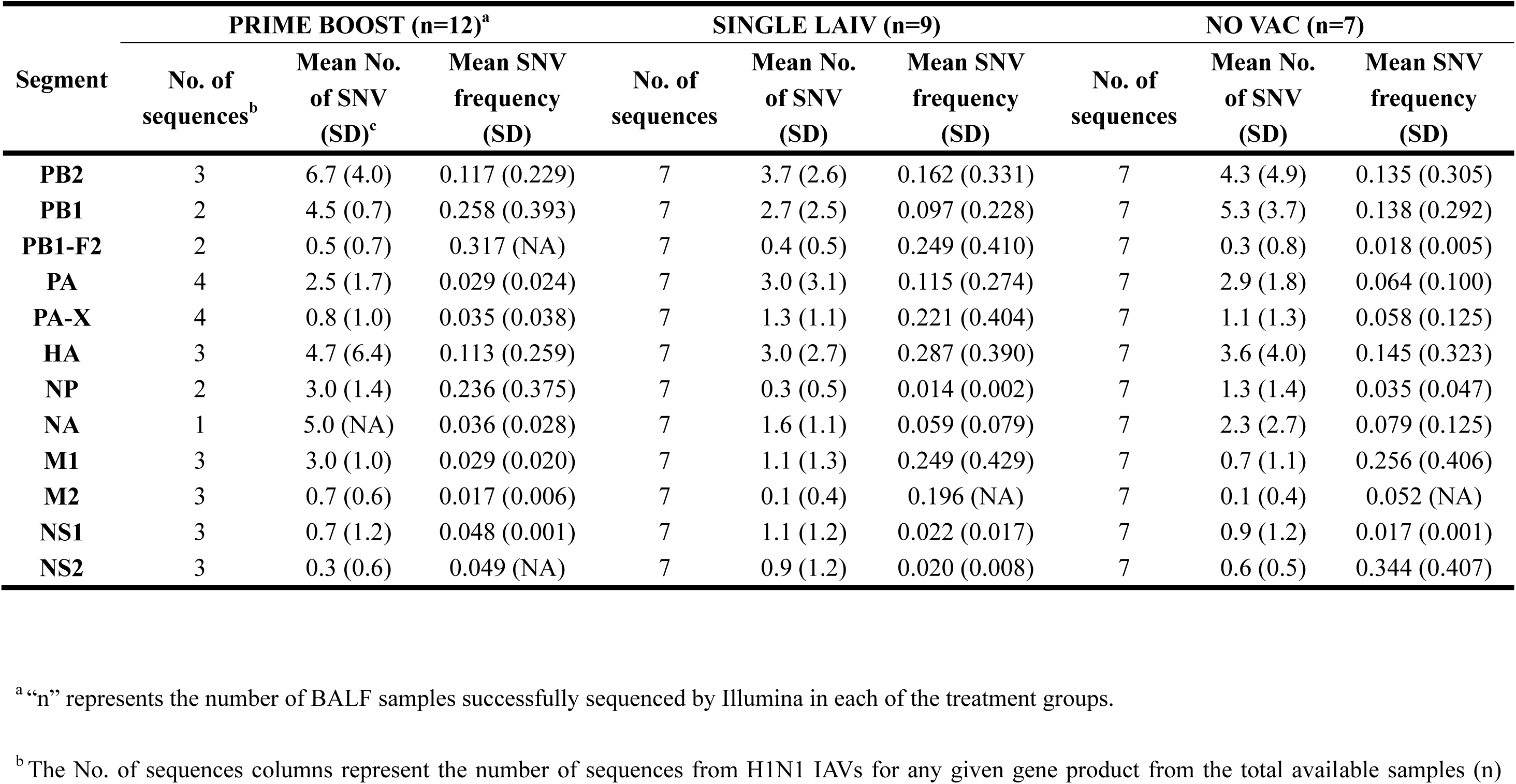
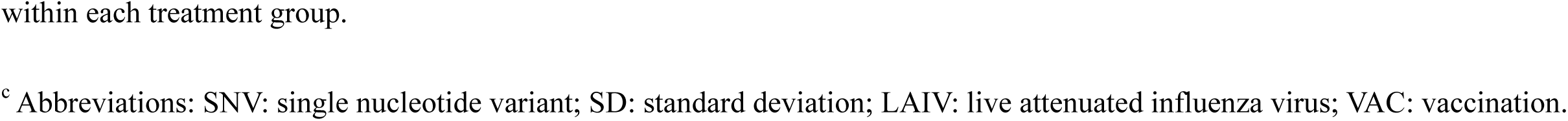
Average number and frequency of single nucleotide variants (SNV) detected on H1N1 sequences of broncho alveolar lavage fluid

| Segment | PRIME BOOST (n=12)<sup>a</sup> |  |  | SINGLE LAIV (n=9) |  |  | NO VAC (n=7) |  |  |
| --- | --- | --- | --- | --- | --- | --- | --- | --- | --- |
|  | No. of sequences<sup>b</sup> | Mean No. of SNV (SD)<sup>c</sup> | Mean SNV frequency (SD) | No. of sequences | Mean No. of SNV (SD) | Mean SNV frequency (SD) | No. of sequences | Mean No. of SNV (SD) | Mean SNV frequency (SD) |
| **PB2** | 3 | 6.7 (4.0) | 0.117 (0.229) | 7 | 3.7 (2.6) | 0.162 (0.331) | 7 | 4.3 (4.9) | 0.135 (0.305) |
| **PB1** | 2 | 4.5 (0.7) | 0.258 (0.393) | 7 | 2.7 (2.5) | 0.097 (0.228) | 7 | 5.3 (3.7) | 0.138 (0.292) |
| **PB1-F2** | 2 | 0.5 (0.7) | 0.317 (NA) | 7 | 0.4 (0.5) | 0.249 (0.410) | 7 | 0.3 (0.8) | 0.018 (0.005) |
| **PA** | 4 | 2.5 (1.7) | 0.029 (0.024) | 7 | 3.0 (3.1) | 0.115 (0.274) | 7 | 2.9 (1.8) | 0.064 (0.100) |
| **PA-X** | 4 | 0.8 (1.0) | 0.035 (0.038) | 7 | 1.3 (1.1) | 0.221 (0.404) | 7 | 1.1 (1.3) | 0.058 (0.125) |
| **HA** | 3 | 4.7 (6.4) | 0.113 (0.259) | 7 | 3.0 (2.7) | 0.287 (0.390) | 7 | 3.6 (4.0) | 0.145 (0.323) |
| **NP** | 2 | 3.0 (1.4) | 0.236 (0.375) | 7 | 0.3 (0.5) | 0.014 (0.002) | 7 | 1.3 (1.4) | 0.035 (0.047) |
| **NA** | 1 | 5.0 (NA) | 0.036 (0.028) | 7 | 1.6 (1.1) | 0.059 (0.079) | 7 | 2.3 (2.7) | 0.079 (0.125) |
| **M1** | 3 | 3.0 (1.0) | 0.029 (0.020) | 7 | 1.1 (1.3) | 0.249 (0.429) | 7 | 0.7 (1.1) | 0.256 (0.406) |
| **M2** | 3 | 0.7 (0.6) | 0.017 (0.006) | 7 | 0.1 (0.4) | 0.196 (NA) | 7 | 0.1 (0.4) | 0.052 (NA) |
| **NS1** | 3 | 0.7 (1.2) | 0.048 (0.001) | 7 | 1.1 (1.2) | 0.022 (0.017) | 7 | 0.9 (1.2) | 0.017 (0.001) |
| **NS2** | 3 | 0.3 (0.6) | 0.049 (NA) | 7 | 0.9 (1.2) | 0.020 (0.008) | 7 | 0.6 (0.5) | 0.344 (0.407) |

<sup>a</sup>“n” represents the number of BALF samples successfully sequenced by Illumina in each of the treatment groups.
<sup>b</sup> The No. of sequences columns represent the number of sequences from H1N1 IAVs for any given gene product from the total available samples (n)
within each treatment group.
<sup>c</sup> Abbreviations: SNV: single nucleotide variant; SD: standard deviation; LAIV: live attenuated influenza virus; VAC: vaccination.

Within the SNVs dataset of the H3N2 virus, we observed 364 SNVs out of 257,699 sequenced consensus nucleotides (205 nonsynonymous, 153 synonymous, 6 stop-gained) in 21 BALF samples (Figure 4 - figure supplement 1B and Figure 4 – source data 2). Similar to H1N1, the H3N2 SNVs were dominated by low-frequency variants (Figure 4E). There were 187 (91.22%) nonsynonymous, 120 (78.43%) synonymous, and 6 (100%) stop-gained SNVs whose frequency was below 10%. About 4.40% (16/364) of H3N2 SNVs were presented at consensus level which was less than the SNVs presented for the H1N1 virus (p=0.007, Chi-square test). We summarized the SNVs quantity and frequency of the H3N2 virus by gene segment and group in Table 3, and no statistical differences were detected in the average SNV number between treatment groups (p = 0.27 – 1.00, Kruskal-Wallis rank-sum test). The differences of SNVs frequencies in each gene segment were not significant between treatment groups, except the frequency of N2 SNVs detected in PRIME BOOST pigs was higher than that of SINGLE LAIV pigs (p=0.046, Dunn’s test with Benjamini-Hochberg correction). Consistent with the H1N1 virus, we did not find a significant impact of immunity on H3 variations (Figures 4F-H). Specifically, there were no nucleotide changes located in the H3 antigenic regions, and we did not detect any associations between the H3 nucleotide divergence and H3 specific HI titers (R=-0.203, p=0.39, Spearman’s Rank-Order Correlation test) or H3 specific IFN-γ secreting cell counts (R=-0.046, p=0.87, Spearman’s Rank-Order Correlation test). The detailed results of H1/H3 HI titers and H1/H3 specific IFN-γ secreting cell counts for each vaccinated and non-vaccinated pig can be found in [32].

**Table 3.**
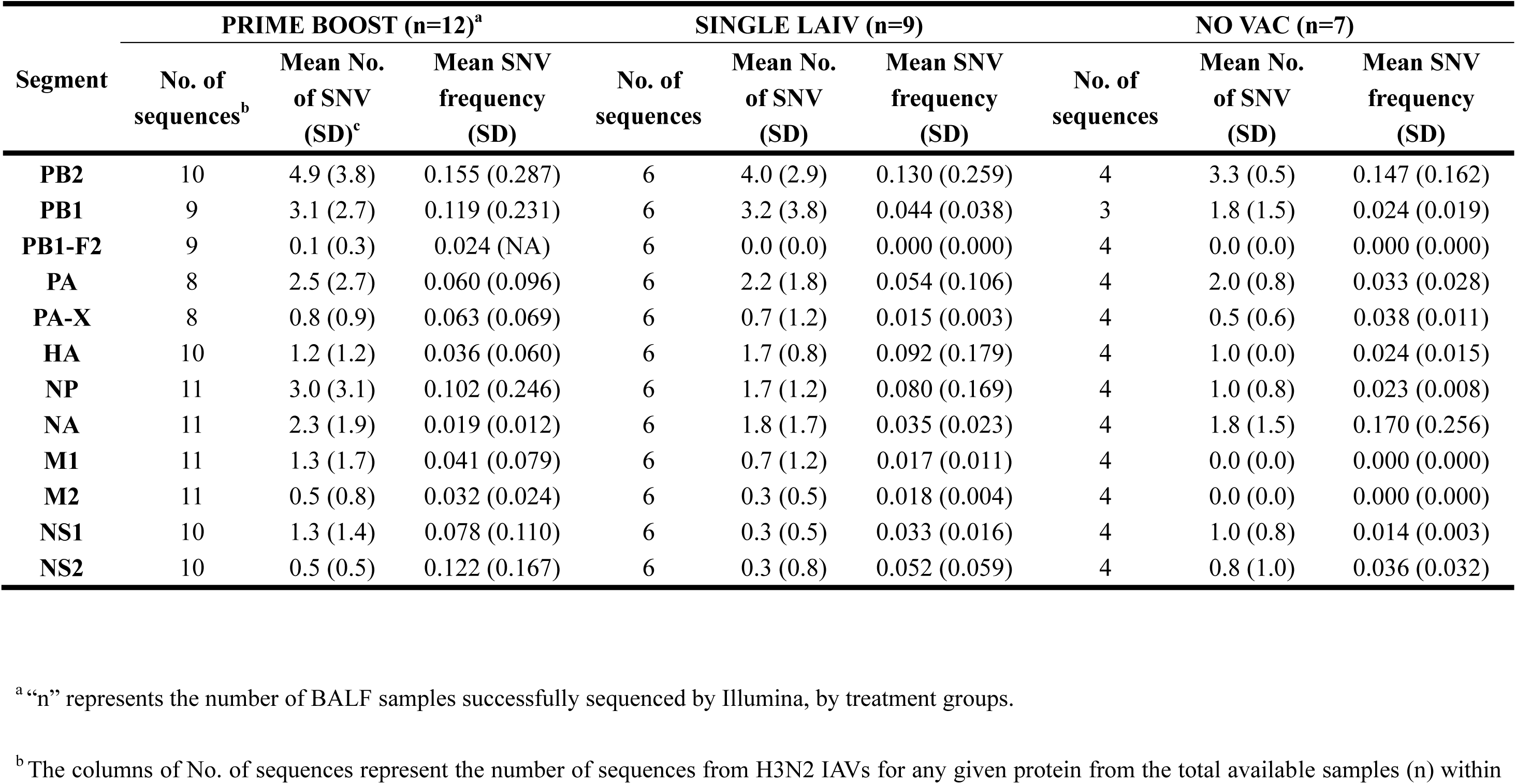
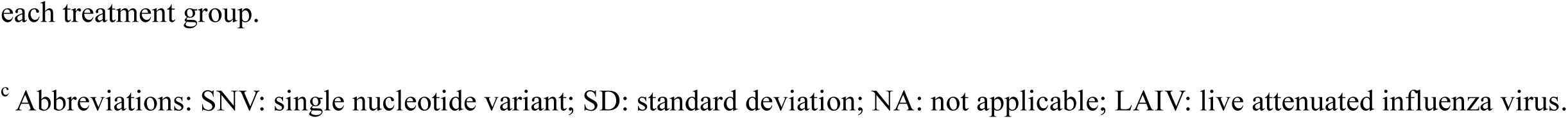
Average number and frequency of single nucleotide variants (SNV) detected on H3N2 sequences of broncho alveolar lavage fluid

| Segment | PRIME BOOST (n=12)<sup>a</sup> |  |  | SINGLE LAIV (n=9) |  |  | NO VAC (n=7) |  |  |
| --- | --- | --- | --- | --- | --- | --- | --- | --- | --- |
|  | No. of sequences<sup>b</sup> | Mean No. of SNV (SD)<sup>c</sup> | Mean SNV frequency (SD) | No. of sequences | Mean No. of SNV (SD) | Mean SNV frequency (SD) | No. of sequences | Mean No. of SNV (SD) | Mean SNV frequency (SD) |
| **PB2** | 10 | 4.9 (3.8) | 0.155 (0.287) | 6 | 4.0 (2.9) | 0.130 (0.259) | 4 | 3.3 (0.5) | 0.147 (0.162) |
| **PB1** | 9 | 3.1 (2.7) | 0.119 (0.231) | 6 | 3.2 (3.8) | 0.044 (0.038) | 3 | 1.8 (1.5) | 0.024 (0.019) |
| **PB1-F2** | 9 | 0.1 (0.3) | 0.024 (NA) | 6 | 0.0 (0.0) | 0.000 (0.000) | 4 | 0.0 (0.0) | 0.000 (0.000) |
| **PA** | 8 | 2.5 (2.7) | 0.060 (0.096) | 6 | 2.2 (1.8) | 0.054 (0.106) | 4 | 2.0 (0.8) | 0.033 (0.028) |
| **PA-X** | 8 | 0.8 (0.9) | 0.063 (0.069) | 6 | 0.7 (1.2) | 0.015 (0.003) | 4 | 0.5 (0.6) | 0.038 (0.011) |
| **HA** | 10 | 1.2 (1.2) | 0.036 (0.060) | 6 | 1.7 (0.8) | 0.092 (0.179) | 4 | 1.0 (0.0) | 0.024 (0.015) |
| **NP** | 11 | 3.0 (3.1) | 0.102 (0.246) | 6 | 1.7 (1.2) | 0.080 (0.169) | 4 | 1.0 (0.8) | 0.023 (0.008) |
| **NA** | 11 | 2.3 (1.9) | 0.019 (0.012) | 6 | 1.8 (1.7) | 0.035 (0.023) | 4 | 1.8 (1.5) | 0.170 (0.256) |
| **M1** | 11 | 1.3 (1.7) | 0.041 (0.079) | 6 | 0.7 (1.2) | 0.017 (0.011) | 4 | 0.0 (0.0) | 0.000 (0.000) |
| **M2** | 11 | 0.5 (0.8) | 0.032 (0.024) | 6 | 0.3 (0.5) | 0.018 (0.004) | 4 | 0.0 (0.0) | 0.000 (0.000) |
| **NS1** | 10 | 1.3 (1.4) | 0.078 (0.110) | 6 | 0.3 (0.5) | 0.033 (0.016) | 4 | 1.0 (0.8) | 0.014 (0.003) |
| **NS2** | 10 | 0.5 (0.5) | 0.122 (0.167) | 6 | 0.3 (0.8) | 0.052 (0.059) | 4 | 0.8 (1.0) | 0.036 (0.032) |

<sup>a</sup>“n” represents the number of BALF samples successfully sequenced by Illumina, by treatment groups.
<sup>b</sup> The columns of No. of sequences represent the number of sequences from H3N2 IAVs for any given protein from the total available samples (n) within
each treatment group.
<sup>c</sup> Abbreviations: SNV: single nucleotide variant; SD: standard deviation; NA: not applicable; LAIV: live attenuated influenza virus.

### Within-pig nucleotide polymorphisms and evolutionary rates of H1N1 and H3N2 viral populations are similar regardless of vaccination status

We calculated the nucleotide diversity (Pi, average number of pairwise nucleotide differences per site) to measure the degree of genetic variation for the H1N1 and H3N2 viral populations within pigs from the different treatment groups (Figure 5A). For both challenge viruses, most coding regions did not have significant differences between pigs from the different groups, except for pigs for the PRIME BOOST group that had significantly higher nucleotide diversity than SINGLE LAIV pigs for NP (p= 0.049, Dunn’s test with Benjamini-Hochberg correction) gene segments of the H1N1 virus. When all coding regions were combined (i.e., whole genome), neither the H1N1 nor the H3N2 viruses from any pig, regardless of treatment group, had statistically significant Pi differences (Figure 5A).

**Figure 5.**
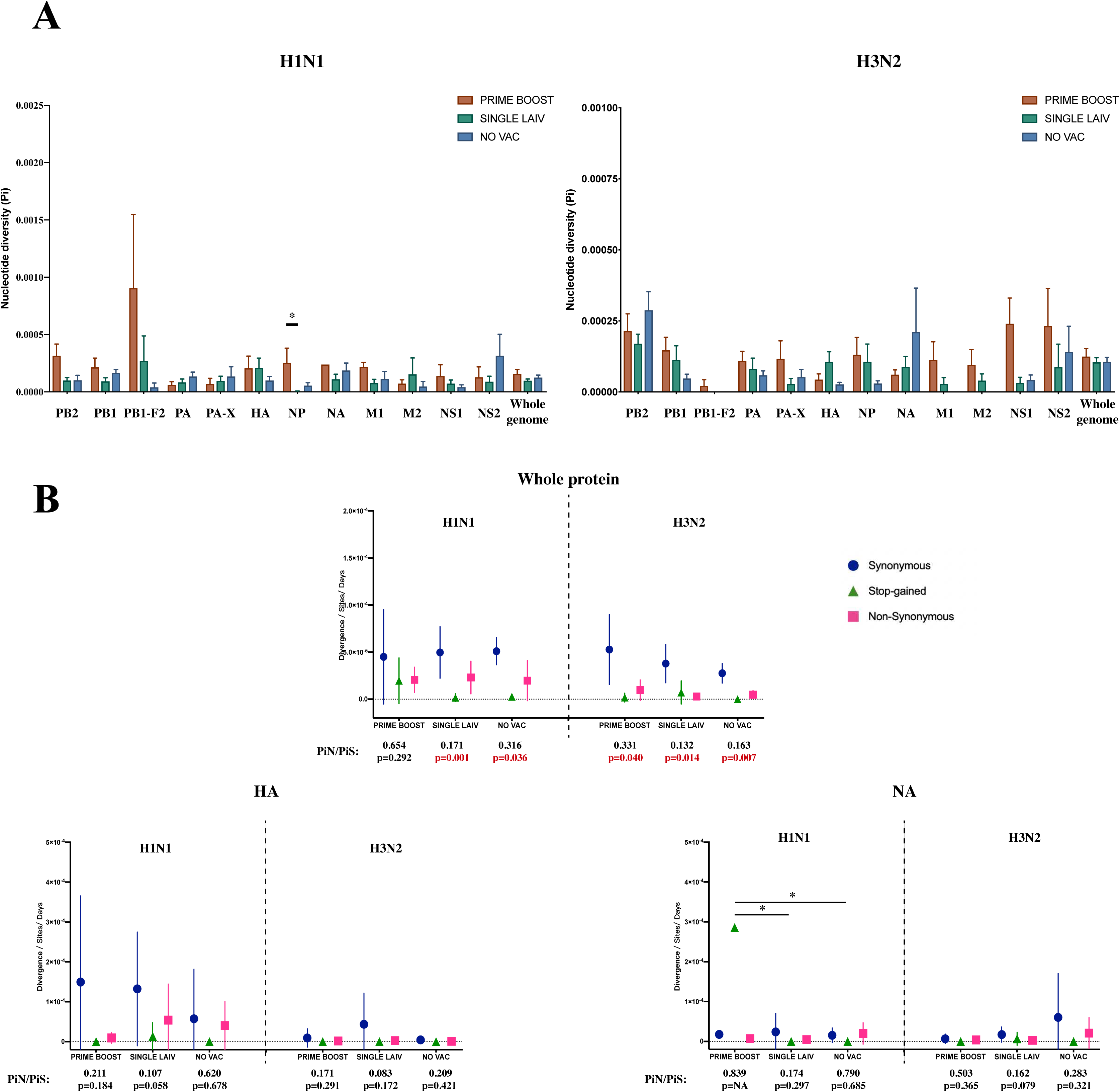
Within-host nucleotide diversity and evolutionary rate of H1N1 and H3N2 influenza A virus (IAV) by coding regions. (A) Nucleotide diversity (Pi) was computed for each coding region of H1N1 and H3N2 viruses for each treatment group. The nucleotide diversity is shown as mean with standard error for PRIME BOOST (brown), SINGLE LAIV (dark green), and NO VAC (dark blue) pigs. The statistical results are noted (* p < 0.05) if the nucleotide diversity of any coding region differed significantly between treatment groups, which were compared by Kruskal-Wallis rank-sum test; the Dunn’s test was used for the multiple pairwise comparisons with Benjamini-Hochberg correction. The standard errors were calculated through 10000 bootstrap resampling with the replacement. (B) The evolutionary rates were calculated separately for H1N1 and H3N2 viruses at synonymous (dark blue circle), nonsynonymous (pink square), and stop-gained (green triangle) sites for each sample on antigenic proteins and whole-genome level. The evolutionary rates are displayed as means with standard deviations and compared by Kruskal-Wallis rank-sum test, followed with the Dunn’s test for the multiple pairwise comparisons; the p-values were corrected by the Benjamini-Hochberg method. The statistical results are noted (* p < 0.05) if the evolutionary rate of any gene and mutational type was significantly different between pigs from any two groups. The piN or piS is the average number of pairwise nonsynonymous or synonymous polymorphism/diversity per nonsynonymous or synonymous site and computed for each gene by the type of mutation. The paired t-test was used to test the null hypothesis that piN = piS, and the significant results (p < 0.05) are marked in red font. The p values are assigned as “NA” if the number of available samples was not enough to perform the statistical analysis and the ratios of piN to piS (piN/piS) are displayed as “NA” if there was no synonymous SNV found in that coding region for pigs in any treatment groups. The piN/piS = 0 indicates no nonsynonymous SNVs were identified in any coding region from pigs in any treatment groups. Figure 5 - figure supplement 1 shows the evolutionary rates and ratios of nonsynonymous to synonymous nucleotide diversity on protein products from H1N1 and H3N2 internal genes.

We calculated the evolutionary rates for H1N1 and H3N2 viruses at the whole genome and at the gene segment level to assess the pace of synonymous, nonsynonymous, and stop-gainedmutations accumulated in the pigs by treatment (Figure 5B and Figure 5 - figure supplement 1). Only the rate of stop-gained mutations in the N1 gene differed between groups (PRIME BOOST – SINGLE LAIV: p = 0.0009, PRIME BOOST - NO VAC: p = 0.0004, Dunn’s test with Benjamini-Hochberg correction). For all other genome changes in the H1N1 and H3N2, similar rates for all three types of mutations between any two individual groups did not differ significantly in any individual IAV coding region within pigs from different vaccination statuses.

The comparison between values of nonsynonymous nucleotide diversity (piN) and synonymous nucleotide diversity (piS) is a measure of within-host virus diversity that can be used to infer the types of selective forces from positive (Darwinian) selection (piN/piS >1), Purified (negative) selection (piN/piS < 1), and genetic drift (piN/piS ∼ 1). The detailed values of piN and piS for each treatment group and coding region are summarized in Supplementary file 2. At the whole-genome level, both H1N1 and H3N2 viruses exhibited significant piN < piS within pigs from all three groups, except for H1N1 in PRIME BOOST pigs (piN/piS = 0.654, p = 0.292, paired t-test), which showed signs of purified selection (Figure 5B). For the majority of gene segments, IAV nucleotide diversity at the gene segment level exhibited piN < piS without showing significant differences for the H1N1 or the H3N2 viruses within pigs regardless of treatment groups. The exceptions for IAV nucleotide diversity at the gene segment level were found for these few within-host instances, e.g., H1N1 PB2 gene in SINGLE LAIV (piN/piS = 0.066, p = 0.021, paired t-test) and NO VAC (piN/piS = 0.291, p = 0.047, paired t-test) pigs, the H3N2 PB2 gene in PRIME BOOST (piN/piS = 0.198, p = 0.036, paired t-test) and SINGLE LAIV (piN/piS = 0.082, p = 0.014, paired t-test) pigs, and H1N1 PB1 gene in NO VAC (piN/piS = 0.223, p = 0.036, paired t-test) pigs (Figure 5 - figure supplement 1). Moreover, we did not find any evidence of positive selection in any coding region for both H1N1 and H3N2 viruses in any of the pigs.

When using a Spearman’s rank correlation coefficient to compute crrelations between the intensity of swine humoral (HA-specific HI titers) and cellular (HA-specific IFN-γ secreting cell counts) immunity versus the IAV HA Pi, PiN, or PiS values of individual pigs from the three treatment groups for both H1N1 and H3N2 viruses, only a weak negative association was found between the H3-specific IFN-γ ELISPOT cell counts and the values of PiN for H3 genes (R = -0.512, p = 0.043, Spearman’s Rank-Order Correlation test). For H1 cellular and humoral immunity and H3 humoral immunity, no correlations were detected (Supplementary file 3).

### SNVs were identified at functional sites in both vaccinated and non-vaccinated pigs

IAV biological properties can be altered significantly with key amino acid changes in coding regions. Out of the 206 nonsynonymous mutations within our H1N1 virus dataset, we found 75 of them located at functional relevant sites using data from the Sequence Feature Variant Types tool in the Influenza Research Database [36]. In addition, 93 out of 205 nonsynonymous SNVs were located at functional relevant sites in the H3N2 genomes. The distribution of these SNVs by functional categories within pigs from different groups is summarized in Supplementary files 4 - 6. We did not detect nonsynonymous SNVs located at sites related to H1 and H3 virus receptor binding or antigenic properties. We found the PRIME BOOST pigs had the highest proportion of functionally relevant SNVs annotated among the three groups for both H1N1 (23/82) and H3N2 (62/211) virus. While the SINGLE LAIV pigs had the lowest percentage of annotated functionally related SNVs for H1N1 (23/135) and H3N2 (19/101) among the three groups. However, the differences in the percentage of H1N1 and H3N2 annotated functional SNVs among treatment groups were not significant statistically (H1N1: p=0.1009, H3N2: p=0.1219, Chi-square test). About 17.8 percent (29/163) of H1N1 SNVs and 23.1 percent (12/52) of H3N2 SNVs in NO VAC pigs were identified as functional SNVs that covered most functional classes, except for the H1N1 SNVs related to cross-species transmission and adaption and H3N2 SNVs associated with virus assembly, budding, and release.

The annotated functionally related SNVs on H1N1 and H3N2 virus were present at low frequencies (mean frequencies with SD for H1N1 virus = 0.095 ± 0.221, mean frequencies with SD for H3N2 virus = 0.048 ± 0.117, p = 0.097, unpaired t-test). Moreover, the frequencies of these annotated SNVs were not significantly different between pigs from different treatment groups for both H1N1 (p = 0.473, Kruskal-Wallis rank-sum test) and H3N2 (p = 0.058, Kruskal-Wallis rank-sum test) viruses.

### Limited shared diversity in H1N1 and H3N2 IAV detected in vaccinated pigs

Vaccine-induced immunity may drive genetic selection within a specific genetic region or site of the IAV genome facilitating the same amino acid changes in multiple pigs even if the pigs are housed in different rooms, which suggest a sign of convergent evolution. There were 13 H1N1 and three H3N2 polymorphic amino acid sites identified in at least two pigs (Figures 6A and 6C). Among them, the H1N1 polymorphic sites - PB2 423, PB1 723, and M1 200 were identified as functionally relevant sites and linked to IAV PB2 cap-binding properties, the binding affinity between IAV PB2 and PB1, and vRNP binding during virus budding, respectively [37–41]. None of the three H3N2 polymorphic sites were identified as functionally related sites. However, most of these polymorphic sites were detected in multiple pigs from the same room, which suggested that the shared amino acid changes observed between pigs were largely due to virus transmission rather than virus mutating independently in the pigs. Therefore, we focused on IAV shared amino acid changes in pigs from different rooms. As a result, we found 0, 1 (PB1 V632A), and 2 (HA A242G and PB2 N562I) amino acid changes of H1N1 viruses in at least two PRIME BOOST, SINGLE LAIV, and NO VAC pigs from different rooms, respectively. For the H3N2 virus, we only identified NA G143R in NO VAC pigs from multiple rooms.

**Figure 6.**
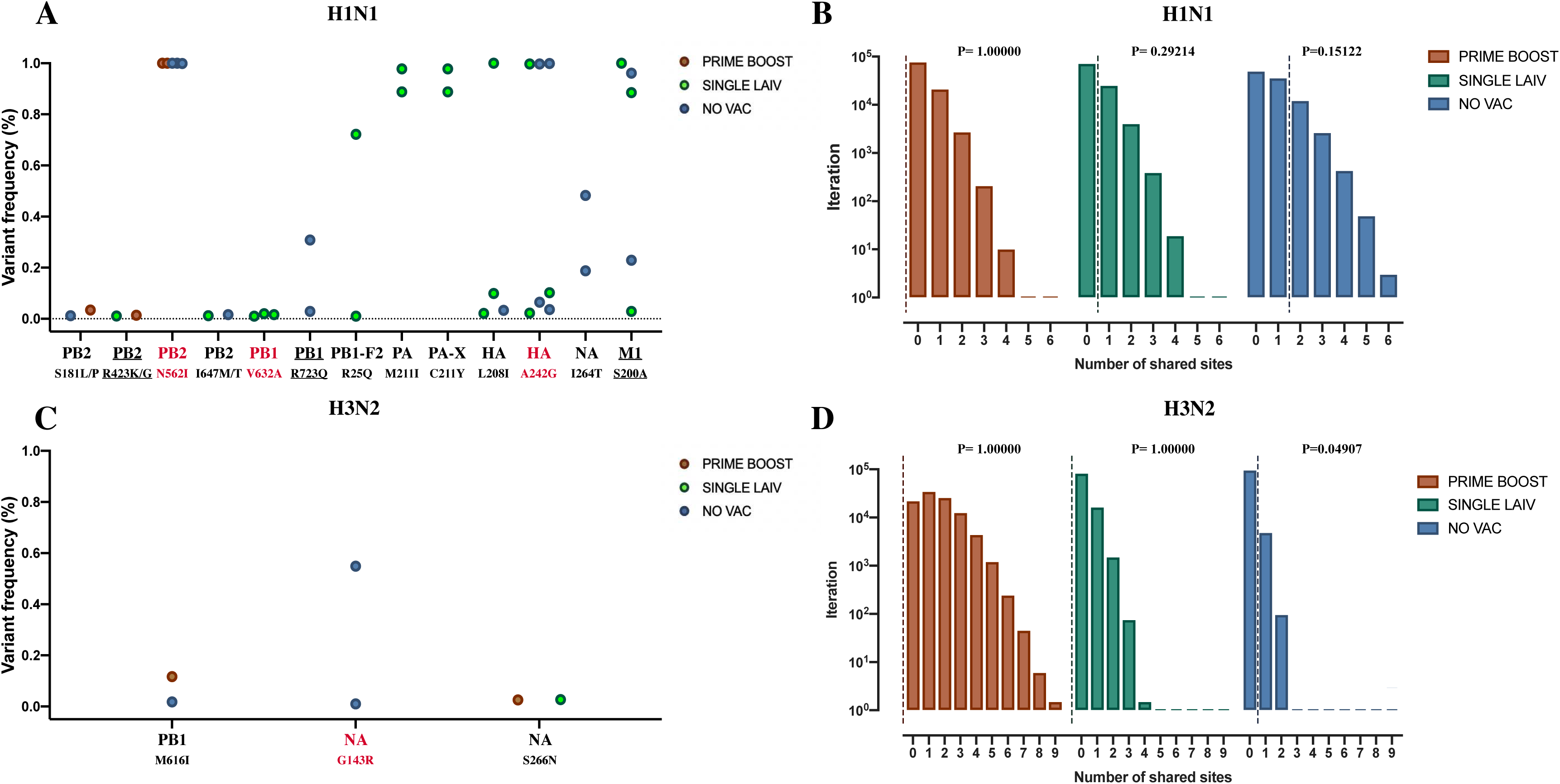
Shared amino acid polymorphisms sites in H1N1 and H3N2 influenza A viruses (IAVs). H1N1 (A) and H3N2 (C) polymorphic amino acid sites detected in at least two samples regardless of treatment groups. The dots represent the same or different amino acid changes at any specific sites and are colored by treatment groups. Sites that were identified as functionally relevant are underlined. The polymorphic sites shared by at least two pigs from different rooms were marked in red. The y-axis indicates the frequency of any amino acid changes within the IAV population. The permutation test of shared amino acid polymorphisms in 70% H1N1 (B) and H3N2 (D) IAV genome in pigs is shown by treatment groups. The x-axis indicates the number of shared polymorphic sites present in at least two samples collected from multiple rooms within the same treatment group. The y-axis indicates the number of iterations that have given number of shared polymorphic sites within each treatment group. The dashed lines represent the actual number of shared amino acid polymorphic sites observed in our data from H1N1 or H3N2 viruses in BALF samples from each group. The same processes of permutation tests on shared amino acid polymorphism sites are also run on the entire genomes of H1N1 and H3N2 IAVs. The results are shown in Figure 6 - figure supplement 1.

We performed the permutation test in 100,000 iterations for each treatment group to evaluate whether the number of shared amino acid sites changes observed in pigs from multiple rooms was more than expected by chance (see Materials and Methods). We found both PRIME BOOST and SINGLE LAIV pigs shared no more polymorphic amino acid sites than expected by chance for H1N1 (p= 1.00000 for PRIME BOOST pigs, p= 0.21631 for SINGLE LAIV pigs) and H3N2 (p= 1.00000 for PRIME BOOST pigs, p= 1.00000 for SINGLE LAIV pigs) viruses. However, the shared diversity in NO VAC pigs was more than random chance for H3N2 viruses (p= 0.03536), but not for H1N1 viruses (p= 0.08396) (Figure 6 - figure supplement 1). The IAV genome is highly constrained, and in a different study, it was shown that about 30% of mutations in IAV genome regions were lethal [42]. The ability of IAV to buffer the fitness effect from the mutations or IAV mutational robustness could profoundly affect the results of IAV shared diversity. Therefore, we assumed that 70% of the IAV genome could tolerate mutations and we ran the permutation test on 70% of each coding region’s amino acid sites (Figures 6B and 6D). The results showed that the shared polymorphisms in pigs of all the three treatment groups were no more than expected by chance for both, H1N1 (p= 1.00000 for PRIME BOOST pigs, p= 0.29214 for SINGLE LAIV pigs, and p= 0.15122 for NO VAC pigs) and H3N2 (p= 1.00000 for PRIME BOOST pigs, p= 1.00000 for SINGLE LAIV pigs) viruses after considering the genome constraint, except the weak genome convergence of H3N2 viruses observed in NO VAC pigs (p = 0.04907). Overall, we observe the minimal affection of vaccination on the degree of IAV convergence in pigs from the different rooms for either the H1N1 or the H3N2 virus.

## Discussion

Understanding the impact of vaccination on IAV within-host evolution is critical for improving overall animal health and productivity while also decreasing the emergence of novel antigenic variants with pandemic potential [12,43,44]. To better mimic field scenarios, we set up an *in vivo* co-infection model where we challenged naïve pigs with either an H1N1 (gamma clade 1A.3.3.2) or an H3N2 (human-like clade 3.2010.1) virus belonging to the dominant subtypes and lineages identified during US pig surveillance [45]. Pigs were co-housed in contact with nonvaccinated pigs or pigs vaccinated by licensed multivalent vaccines following multiple vaccination protocols [32]. The co-infection model allowed us to simultaneously assess the IAV genomic variation and reassortment under immune pressure in pigs and provided the rare view of the full impact of vaccination on IAV evolution. The dataset includes pigs with different levels of virus shedding and the immune responses obtained enable us to define the factors that affect IAV evolution [32]. We directly sequenced and performed the isolation of viral plaques on selected BALF samples without additional virus propagation in cells to ensure that no mutation or reassortants were developed during cell culture. Our results suggested that the pigs receiving two doses of vaccines (PRIME BOOST) had a lower proportion of IAV reassortants with fewer shared amino acid changes when compared to nonvaccinated pigs. A significant number of mutations with abundant functional relevant amino acid changes were present in H1N1 and H3N2 viral populations within pigs regardless of vaccination status. However, we did not detect a major effect of vaccination on IAV within-host diversity as there were few antigenic variants detected in the IAV populations, and we found no differences in the number and frequency of identified SNVs, the evolutionary rates, nor in the nucleotide diversity of IAV in pigs of the different treatment groups.

Within the seven days of the study period, we identified a large number of IAV reassortants with multiple distinct genotypes in the pigs that had been co-infected with the H1 and H3 viruses. This finding is particularly notable when considering the complex ecosystem and management practices implemented in swine farms, and the pigs with various vaccination and infection statuses present in the farms [37–39]. The fluidity of ages and immune status of the pigs in swine farms enable viruses to continually infect susceptible pigs and create an ideal breeding ground for IAV to reassort and circulate [40]. Given the background that the co-existence of viruses with distinct lineages and genotypes is common in pig populations, the frequency of IAV reassortment could be extensive, especially considering that most of the pigs need to be housed approximately for 24–26 weeks in farms [12]. Our study provides initial evidence that vaccination can play a role in decreasing the generation and emergence of new reassortant viruses in pigs. These results, if confirmed in field conditions, show the significance of IAV vaccination in pig populations not only by restricting the virus circulation but also by reducing the IAV genotypic diversity which is important for the overall IAV control at the human-animal interface [46].

The factors driving reassortment are complicated, and for IAV reassortment to occur two parental viruses need to reach multiple criteria [47]. The co-infection of distinct IAV viruses at the cellular level is an essential prerequisite for IAV reassortment [48]. Numerous factors affecting IAV cellular co-infections have been elucidated in vitro or under experimental conditions, including the IAV challenge dose, the time interval between primary and secondary infection, location of initial virus replication, and tissue tropism of IAV viruses [49,50]. Our study found that pigs co-infected for a longer duration tended to generate more reassortants, which showed that vaccination could minimize IAV reassortment by reducing the possibility of pigs being co-infected by distinct viruses. It has been shown that most IAV reassortants emerge with lower fitness compared with parental viruses, and evidence suggests that the strength of negative selection on the reassortants is positively associated with the genetic distance between their parental strains [51]. In our study, we observed that the IAV internal genes were exchanged fairly easily between the two challenge, i.e., parental, viruses compared to the antigenic genes. Considering the nucleotide homology of internal genes between the challenge viruses was much higher (over 90%) than that of the antigenic genes (∼53%), the mismatch of package signals and functional proteins could explain these results given that these restraining factors facilitate virus reassortment between similar strains rather than in distantly related viruses [47,50,52].

The high plasticity of the IAV genome and the fast-expanding nature of IAV favors the generation of progeny virions with slightly different mutation “signatures” [5]. Although most progeny variants have a deleterious effect on the overall fitness of the viral populations, some of the beneficial mutants may go through positive selection and can quickly dominate after sudden changes in the environment thereby shifting the fitness landscape, which may maximize the possibility of IAV to replicate in harsh conditions [53]. As a result, IAV may increase mutation rates at the cost of fitness to enrich the genetic pool and accumulate more beneficial variants with high frequencies favored by positive selection [54,55]. In a stable environment, gaining fitness is a priority for IAV rather than the accumulation of genetic variations at the consensus level [53]. Therefore, purified selection takes place as the major selective force that continually removes new variants and generates IAV populations with a high proportion of low-frequency variants [35]. Previous studies have identified that host immunity could drive positive selection on the IAV antigenic proteins in the long term [56–58]. However, in our study, we rarely observed the inference of the same selection pressures at any coding regions for the H1N1 and H3N2 viruses, which is consistent with the results of within-host diversity on H5N1 virus [35]. Instead, the within-host diversity of both subtypes was dominated by low-frequency variants and exhibited similar or higher synonymous polymorphisms compared to the nonsynonymous polymorphisms. The variation patterns of IAV suggested that the viral populations may be broadly shaped by purifying selection and genetic drift within pigs regardless of vaccination statuses. In addition to the close mutational spectra we observed on H1N1 and H3N2 viruses in pigs from various treatment groups, we detected limited shared diversity, which developed independently in at least two PRIME BOOST, SINGLE LAIV, or NO VAC pigs for both viruses. Our observation suggested that the vaccination did not drive the progress of IAV genome convergence, at least in the swine lower respiratory tract, when we measured the virus shared diversity between pigs, which in turn reflected the limited selective forces raised by vaccination on viral within-host diversity.

The HA protein is the major target of the host adaptive immune response and crucial to IAV evolution. Within our dataset, there were no amino acid changes that fell in the H1 and H3 antigenic or receptor binding sites. Furthermore, the numbers of SNVs on the HA segment did not correlate with the intensity of humoral and cellular immune responses induced by vaccination. These observations are concordant with the results from Debbink et al. [59], which showed the limited affection of vaccination on antigenic diversity of H3N2 viruses in humans. However, other factors like host type, IAV strain, infection dose, challenge/infection model, virus passage among animals, and observation period may cause discrepancies in the results of selection of the antigenic variants between the studies that explore the influence of immune pressure on IAV within-host diversity [25–27,59]. Taken together, we did not find evidence that vaccination influenced IAV genetic variation in the lower respiratory tract of pigs. The observation of our study aligned with the data from Murcia et al. [25], which suggested that vaccine-induced immunity had minimal impact on the genetic variation of IAV populations in both upper and lower respiratory tracts of pigs.

Evaluating IAV evolution in vaccinated hosts is often difficult to perform. One challenge is to obtain enough qualifying samples in pigs that are well-protected by vaccination since there may be low levels of, or even no, replicating virus. The PRIME BOOST samples analyzed in this study included samples from pigs receiving multiple prime-boost vaccination protocols. However, this did not seem to cause significant differences between groups since they induced a similar HA-specific adaptive immune response and exhibited a comparable level of protective efficacy on pigs against the challenge viruses [32]. Different vaccine combinations may introduce variability in the mutational landscape among PRIME BOOST pigs. However, it was not the study’s primary aim to detect IAV SNVs specific to any vaccination protocol but rather to compare the impact of vaccination on the overall evolutionary trends of IAV in pigs. We also had a limited number of BALF samples available for plaque purification and the number of plaques isolated from each sample was not equal. Although we acknowledge this as a limitation, the overall conclusions were supported by the statistical methods employed to assess the relationship between percent of reassortants and treatment group, which accounted for the impact of sample quantity and the variability of total plaques isolated from each sample.

Understanding the association between IAV vaccination and evolution is hard but necessary. IAV within-host evolution plays a vital role in disease outcomes since mutations and reassortment events can significantly influence the biological characteristics of IAV. In this study, we conclude that pig vaccination should be explored as a way to reduce the generation and emergence of reassortant viruses, and this may be particularly important in farm animals where large populations are housed together. In contrast, we did not observe the same significant effect of vaccination on IAV genetic variation. Our research results provide insights into the complexity of IAV evolution in pigs and will help develop more effective influenza control programs to mitigate the burden of IAV infections in pigs while decreasing the risk of zoonotic infections and preserving public health. We believe our study assembles a comprehensive recognition of IAV evolutionary strategies under immune pressure but does not fully reflect the situation in the field. Therefore, more studies are needed to verify our findings under field conditions, which is essential for developing IAV vaccines and surveillance strategies.

## Materials and Methods

### Vaccine-challenge experiments in pigs

All the BALF samples analyzed in this study were obtained from pigs infected using a co-infection seeder pig model using an H1N1 and an H3N2 strain to evaluate the protective efficacy of different prime-boost vaccination protocols in pigs [32]. All the animal work followed the protocols approved by the University of Minnesota Institutional Animal Care and Use Committee (Protocol ID: 1712-35407A) and the Institutional Biosafety Committee (Protocol ID: 1508-32918H).

The infection seeder pig model was set up to mimic field conditions were pigs become infected by contact transmission by challenging 14 eight-week-old unvaccinated pigs with either H1N1 (A/swine/Minnesota/PAH-618/2011) or H3N2 (A/swine/Minnesota/080470/2015) IAV strains and commingling them 2 days post challenge with the vaccinated pigs. The homology of nucleotides and amino acids between the H1N1 and H3N2 challenge viruses are summarized in Supplementary file 7. Whole genome sequences of the two challenge viruses have been deposited in Genbank with accession numbers from MT377710 to MT377725. The H1N1 and H3N2 seeder pigs were evenly distributed as pairs of two in each room and served as the infection source of H1 and H3 viruses to pigs from different treatment groups. The vaccines used for different vaccination protocols included an inactivated commercial quadrivalent vaccine (COM), an inactivated autogenous trivalent vaccine (AUT), and a live attenuated bivalent vaccine (LAIV). The detailed information for the vaccines used in this study can be found in [32]. Seventy pigs were randomly assigned to seven treatment groups with different vaccine combinations which included four whole inactivated vaccinated (WIV) groups (COM/COM, AUT/AUT, AUT/COM, and COM/AUT), two live attenuated vaccinated (LAIV) groups (LAIV/NONE and LAIV/COM) and one positive unvaccinated control group (NO VAC/CHA). No samples from AUT/COM pigs were used in the current study since pigs were protected by vaccination and no IAV positive samples were obtained. Pigs were primed with the first vaccine administration at approximately three weeks of age, boosted at six weeks of age, commingled with seeder pigs at eight weeks of age, and euthanized at nine weeks of age. Pigs vaccinated with inactivated vaccines (COM or AUT) and the positive control groups were housed together in the same rooms and evenly distributed in 5 rooms. Pigs receiving the LAIV treatment were evenly distributed in 2 rooms. Each room contained two seeder and ten treatment pigs for a total of 12 pigs per room. Nasal swabs were collected daily for all the pigs from 2 to 6 days post contact. BALF samples were taken at necropsy at seven days post contact with the seeder pigs. HA subtyping real-time PCR was performed in all the nasal swabs and BALF samples using the VetMAX gold SIV subtyping kit (Life Technologies, Austin, TX, USA). The methods and results for testing the BALF samples by IAV matrix gene real-time RT-PCR, virus titration, IFN-γ ELISPOT cell counts on the lymph nodes and hemagglutinin inhibition (HI) titers for the serum obtained from each pig are shown in [32]. The BALF samples with Ct values below 35 obtained using the IAV matrix gene real-time RT-PCR were selected for whole-genome sequencing directly from the sample, without any virus propagation, to identify the IAV within-host diversity in pigs.

To evaluate IAV reassortment happening in naïve and vaccinated pigs, we selected pigs from 2 rooms housing pigs vaccinated with inactivated vaccines and one room housing LAIV vaccinated pigs. Seeder pigs in these rooms shed relatively high quantity of both challenge viruses (Ct values ranging between 21.15 and 30.51 when contacted with vaccinated pigs). BALF samples from pigs that yielded virus replication by virus titration (TCID50≥1.75/ml) were used for viral plaque purification. Information of BALF samples available for direct sequencing and plaque assay is summarized in Table 1. For analysis purposes, pigs were grouped in three groups named a) prime boost (COM/COM, AUT/AUT, COM/AUT and LAIV/COM pigs), b) single LAIV (LAIV/NONE pigs) and c) non-vaccinated (NO VAC/CHA pigs) groups.

### Plaque library preparation

Plaque assay to purify individual virions was performed on Madin-Darby Canine Kidney (MDCK) cell monolayers [30]. Briefly, BALF samples were 10-fold serially diluted using IAV growth medium, which made up of Dulbecco’s modified Eagle medium (DMEM, Gibco Invitrogen, Carlsbad, CA, USA), 4% BSA fraction V 7.5% solution (Gibco-Life Technologies, Carlsbad, California, USA), 0.15% 1-mg/ml TPCK trypsin (Sigma-Aldrich, St. Louis, USA), and 1% antibiotic-antimycotic (Gibco, Life Technologies, New York, NY, USA). MDCK cells were obtained from the University of Minnesota Veterinary Diagnostic Laboratory (VDL) and cultured in six-well plates were washed twice using Hanks’ Balanced Salt Solution (HBSS, BioWhittaker, Verviers, Belgium) with 0.15% 1-mg/ml TPCK trypsin (Sigma-Aldrich, St. Louis, USA), then inoculated using the diluted BALF samples and incubated for one hour in 37 ℃ at 5% CO2 incubator. After melting in 70 ℃ water bath, one volume of 4% Agarose Gel (Gibco, Life Technologies) was mixed by a three-volume of preheated IAV growth media to make the overlay gel liquid and kept at 37 ℃ water bath. After one-hour incubation, the diluted samples were aspirated and gently replaced by the warm overlay gel liquid at room temperature. When the overlayed gel became solid, the plates were invertedly incubated in a CO2 incubator for 3 to 5 days. Up to 30 visualized plaques were randomly picked up from each sample using micropipette tips and propagated individually on MDCK cells. The isolated plaques were stored at -80 ℃ for further whole-genome sequencing. The numbers of plaques isolated from each BALF sample are summarized in Supplementary file 2.

### RNA extraction, next generation sequencing, and quality control

RNA extraction from BALF samples of plaque isolates was performed using the MagMax Viral RNA isolation kit (Ambion, Life Technologies, USA) [60]. The viral cDNA was amplified from extracted RNA through one-step reverse transcription-PCR amplification by using SuperScript III One-Step RT-PCR system with High Fidelity Platinum Taq DNA Polymerase (Invitrogen, Life Technologies, USA) with degenerate primers (10uM MBTuni-12M and MBTuni-13) [61]. The PCR product was checked by NanoDrop 1000 (Thermo Fisher Scientific) and cleaned up by the Qiagen QIAquick PCR Purification Kit (QIAGEN, USA). Purified PCR products were fragmented and tagged with the indexed adaptors using the Nextera DNA XT Sample Preparation Kit (Illumina, San Diego, CA, USA). The sequence library was quantified by using the Quant-iT PicoGreen dsDNA Assay Kit (Invitrogen). The barcoded library was pooled in equimolar concentrations and multiplexed and sequenced by 150 bp paired-ends on Illumina platform at the University of Minnesota Genomics Center (UMGC). The sequence data obtained from UMGC was analyzed using the software available at the University of Minnesota Supercomputing Institute (MSI) platform. The raw reads were assessed for sequence quality using Fast-QC and trimmed by Trimmomatic [62,63]. The trimming on the BALF sequences was conducted using the command: java -jar path-to-trimmomatic-0.33.jar PE input.fastq output.fastq ILLUMINACLIP: path-to-adaptor.fasta:2:30:10 LEADING:3 TRAILING:3 SLIDINGWINDOW:5:30 MINLEN:100. The command removes the adaptors and the bases with Q-score below 3 at the beginning and end of the raw reads, trimming the sequences in sliding windows of 5 base pairs and cutting the scan bases with the average Q-scores under 30. The reads were discarded if trimmed to the length below 100 bps. The trimming of the plaque sequences was performed using the same command but with different parameters: java -jar path-to-trimmomatic-0.33.jar PE input.fastq output.fastq ILLUMINACLIP: path-to-adaptor.fasta:2:30:10 LEADING:3 TRAILING:3 SLIDINGWINDOW:4:20 MINLEN:36, which trimmed the raw reads with the sliding windows of 4 base pairs and cutting the scan bases with the average Q-scores under 20. Reads were omitted if the length was below 36 after trimming.

### Reassortant recognition of viral plaques

The trimmed reads of viral plaque isolates were de-novo assembled by Shovill [64]. The assembled contigs were annotated by OctoFLU for the initial inspection and sorted by IAV genes [65]. Within each sample, the same IAV gene’s contigs were omitted if the sequence length was < 70% of the open reading frame (ORF) or < 20% of the overall k-mer coverage on corresponding IAV segments. Occasionally, the longest contigs (∼ 1.5% of total analyzed sequences) of any plaque isolates were preserved for genotyping if no sequences covered 70% of ORF for any of their IAV segments. The enrolled consensus sequences from isolated plaques for IAV reassortant identifications were shown in Figure 2 – source data 2, and aligned with the sequences from the curated reference package from OctoFLU, and the sequences of H1N1 and H3N2 challenge viruses were aligned using the MUSCLE program in Geneious (version 2021.0.3) [65–67]. The plaque gene segments’ sources were initially checked by comparing the genetic distance between plaque sequences with the reference sequences of H1N1 and H3N2 challenge viruses in Geneious (version 2021.0.3). IQ-TREE was used to construct the maximum likelihood tree of each gene segment in 1000 bootstrap replicates with the best-fit nucleotide substitution model auto-detected by its curate package – ModelFinder to further verify the origin of plaque segments [68]. The tree files were visualized in FigTree [69]. Based on the origins of the eight gene segments, a genotype was assigned to each plaque isolate. A viral plaque was considered a reassortant if there were one or more gene segment substitutions of both parental H1N1 and H3N2 challenge viruses. A mixed genotype denomination was assigned to those reassortants from plaques that in a given gene segment, had complete gene segments from both of the parental viral strains. The maximum likelihood tree for the sequences of each gene segment is displayed in Figure 2 – source data 1.

### Identification of within-host variants

The variant calling pipeline has been described previously elsewhere [35]. In this study, we only identified the within-host variants on the gene reads directly sequenced from the BALF samples. Briefly, the trimmed reads were imported and mapped to the reference sequences of the H1N1 and H3N2 challenge viruses in Geneious (version 2021.0.3) [66]. The mapped reads were exported in SAM format, sorted by Picard Sortsam (command: java -jar picard.jar SortSam INPUT=input_file.sam OUTPUT=output_file.sam SORT_ORDER=coordinate), and duplicate reads removed using Picard MarkDuplicates (command: java -jar picard.jar MarkDuplicates REMOVE_DUPLICATES=true INPUT=input_file.sam OUTPUT=output_file.sam METRICS_FILE=test.bam.metrics) to avoid PCR bias (http://broadinstitute.github.io/picard/). The files were converted to mpileup files by Samtools (https://github.com/samtools/samtools) and the viral variants were called by VarScan (https://github.com/dkoboldt/varscan) with the command : samtools mpileup -f reference_file.fasta input_file.sam | java -jar VarScan.v2.3.9.jar pileup2snp > output_file.vcf [70,71]. The reported variants were filtered with a minimum depth of 100 reads, the minimum frequency of 1%, mean PHRED quality score of 30, and with the variant detected in both forward and reverse reads by performing the command: java -jar VarScan.v2.3.9.jar filter input_file.vcf --min-coverage 100 --min-avg-qual 30 --min-var-freq 0.01 --output-vcf 1 > output_file.vcf. For the samples that contained both challenge viruses, the generated variant report was checked and corrected using a custom Python script to ensure no variants were mistakenly recorded in the report due to nucleotide differences between the challenge viruses, since mis-mapping of the reads against the reference templates could occur due to the close genetic distance of internal gene segments between H1N1 and H3N2 viruses (https://github.com/TorremorellLabUMN/Swine-IAV-within-host-evolution/tree/Main/Script/False_SNV_remove). For the called SNVs that located in the same nucleotide sites of H1N1 and H3N2 genomes from the same sample, we will check the original mapping reads and withdraw the false positive SNVs from the variant report. The final identified variants listed in vcf files were parsed and annotated based on their affection of amino acid changes on reference sequences by the custom Python script (https://github.com/TorremorellLabUMN/Swine-IAV-within-host-evolution/tree/Main/Script/SNV_annotation).

### Evolutionary analysis

The within-host evolutionary rates were calculated separately for synonymous, non-synonymous and stop-gained mutations at the whole-genome and individual protein level based on methods described previously [72]. The evolutionary rate was calculated for each sample by summing the frequencies of single nucleotide variants (SNVs) by their mutation type in each protein and divided by available sites and the number of days post infection. All BALF samples were collected at 7 days post-infection, and the available sites for synonymous, non-synonymous, and stop-gained mutations were normalized by multiplying the total length of nucleotide sequences for each protein by 72%, 25%, and 3%, respectively. These proportions of sites in IAV genome available for different mutation types were calculated in previous published research [72], based on counting the proportion of available sites for each mutation type on the genome of A/Victoria/361/2011 (H3N2) by the Nei and Gojobori method (stop-gained mutations are split from the nonsynonymous mutations) [73], and assuming the frequency of transitions is threefold higher than transversions [74–76]. The efficacy of selection for the IAV populations was estimated by calculating the nucleotide diversity (Pi), synonymous nucleotide diversity (piS), and non-synonymous nucleotide diversity (piN) at the protein level based on the identified within-host variants for each sample by SNPGenie [77] (https://github.com/chasewnelson/SNPGenie).

To identify the variants located in functional relevant sites, the data of all currently available functional annotations in H1, H3, N1, N2 and the other internal segments were downloaded from the Sequence Feature Variant Types tool available in the Influenza Research Database [36]. The non-synonymous variants were annotated if they fell into the annotated region or sites. The annotated functional relevant variants were further categorized based on antiviral drug resistance, determinant of pathogenicity, virulence and disease progression, host-virus interaction machinery, virus assembly, budding and release; viral genome transportation, transcription and replication; viral genome/protein interaction and cross-species transmission and adaption.

The permutation test on shared variant sites was performed at the amino acid level to identify whether the BALF samples from each treatment group shared more polymorphism sites than random chance by the custom Python script adapted from https://github.com/blab/h5n1-cambodia/blob/master/figures/figure-5b-shared-sites-permutation-test.ipynb. The permutation test was performed as described before [35]. Briefly, for each treatment group, we counted the number of variable amino acid sites, n, on each coding region for each sample. We started the permutation test for each group by randomly simulating n variable amino acid sites at each coding region for each sample. Within each iteration, we count the total number of shared polymorphic amino acid sites from all IAV coding regions for each group after removing the amino acid sites shared between pigs from the same room. The null distribution was generated by calculating the total number of polymorphic amino acid sites shared by at least two pigs from different rooms for each treatment group through 100,000 simulations of each gene segment and BALF sample. The p-value was calculated for each treatment group as the total number of iterations in which there were shared equal or larger number of polymorphic sites than observed in actual data, divided by the number of simulations which were 100,000. All the Python scripts used for evolutionary analysis are available at https://github.com/TorremorellLabUMN/Swine-IAV-within-host-evolution/tree/Main/Script.

### Statistical analysis

Statistical analysis were conducted with the R program version 3.6.2 [78]. The percentage of IAV reassortants by treatment groups or vaccination status was compared using a binomial logistic regression model, allowing for overdispersion. The pairwise differences between groups were compared using the chi-squared deviance test and the p-value was adjusted for multiple comparisons using the Bonferroni-Holm method. Kruskal-Wallis rank sum test was utilized to compare the means of SNV quantity and frequency, nucleotide diversity (Pi), and evolutionary rates by synonymous mutation, non-synonymous mutation, and stop-gained between treatment groups. The Dunn’s test was performed for the pairwise comparisons, the p values were adjusted using the Benjamini-Hochberg method. The paired t-test was applied to assess the significant differences between the mean piNs and mean piSs within each treatment group for each gene segment. The Spearman’s rank-order correlation test was performed to evaluate the strength and direction of associations between the percentage of IAV reassortants and co-infection days in pigs. The same statistical analysis also was computed to compare the correlation between HI titers (log2 transformed) and ELISPOT cell counts versus the values of nucleotide divergence, Pi, PiN, and PiS of the HA segment in H1N1 and H3N2 viruses.

### Data availability

The raw sequence reads generated in this study have been deposited in SRA (NCBI) under Bioproject accession number PRJNA813974. All the raw datasets and custom python scripts generated in this study are available in the GitHub repository: https://github.com/TorremorellLabUMN/Swine-IAV-within-host-evolution.

## Acknowledgements

This study is supported by the funding from the Zoetis. The authors are gratefully acknowledged the Aaron Rendahl for his help on statistical methodology.

## Figure Supplements

**Figure 2 – figure supplement 1.**
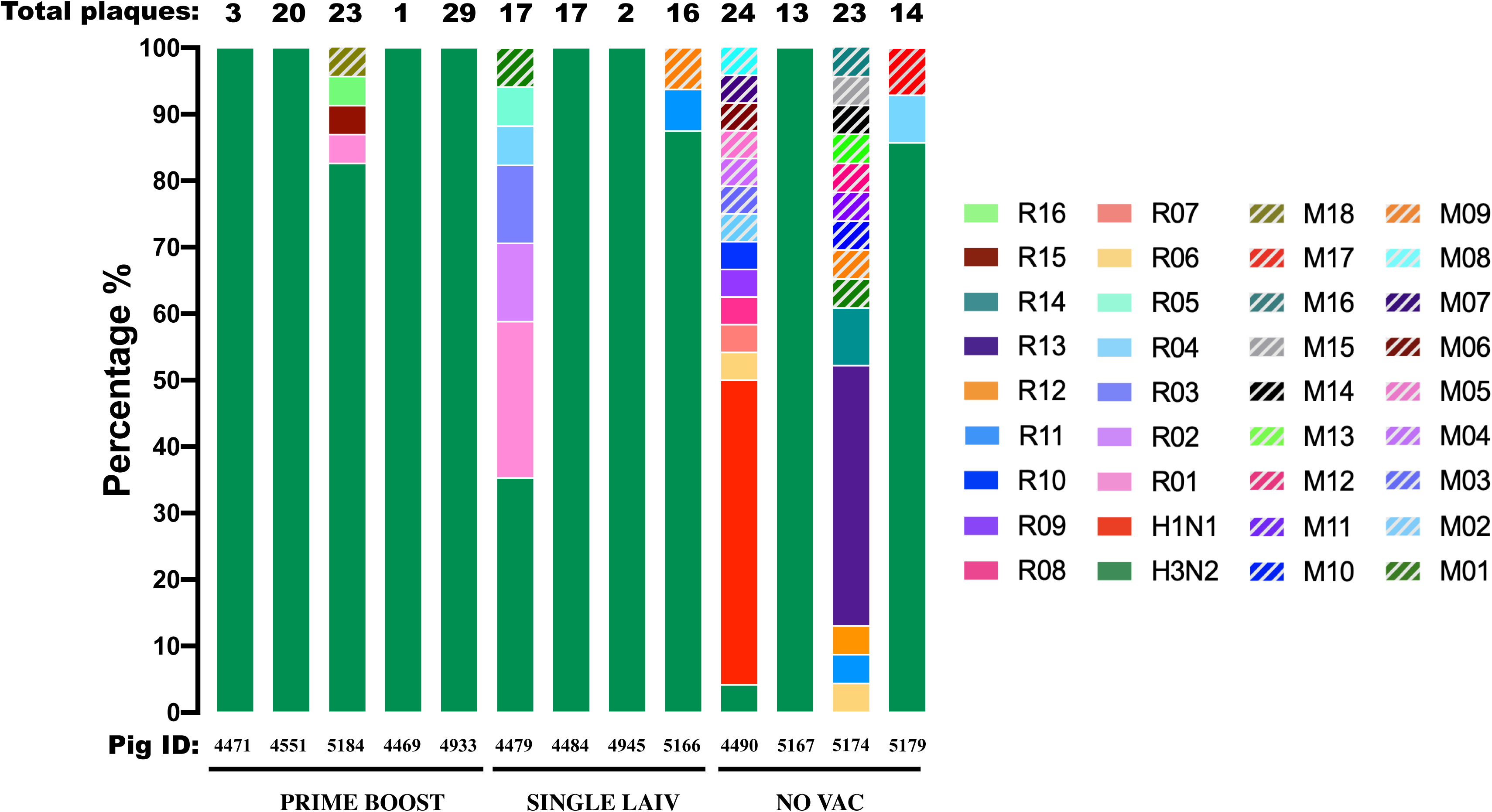
Percentage of reassortant plaques by genotype detected in individual pigs. Each reassortant genotype is shown in a different color except for red and green that show the H1N1 and H3N2 genotypes from the challenge viruses. The number of plaques isolated from individual pigs is displayed above the bar, and the corresponding pig IDs and treatment groups are indicated below the bar.

**Figure 4 - figure supplement 1.**
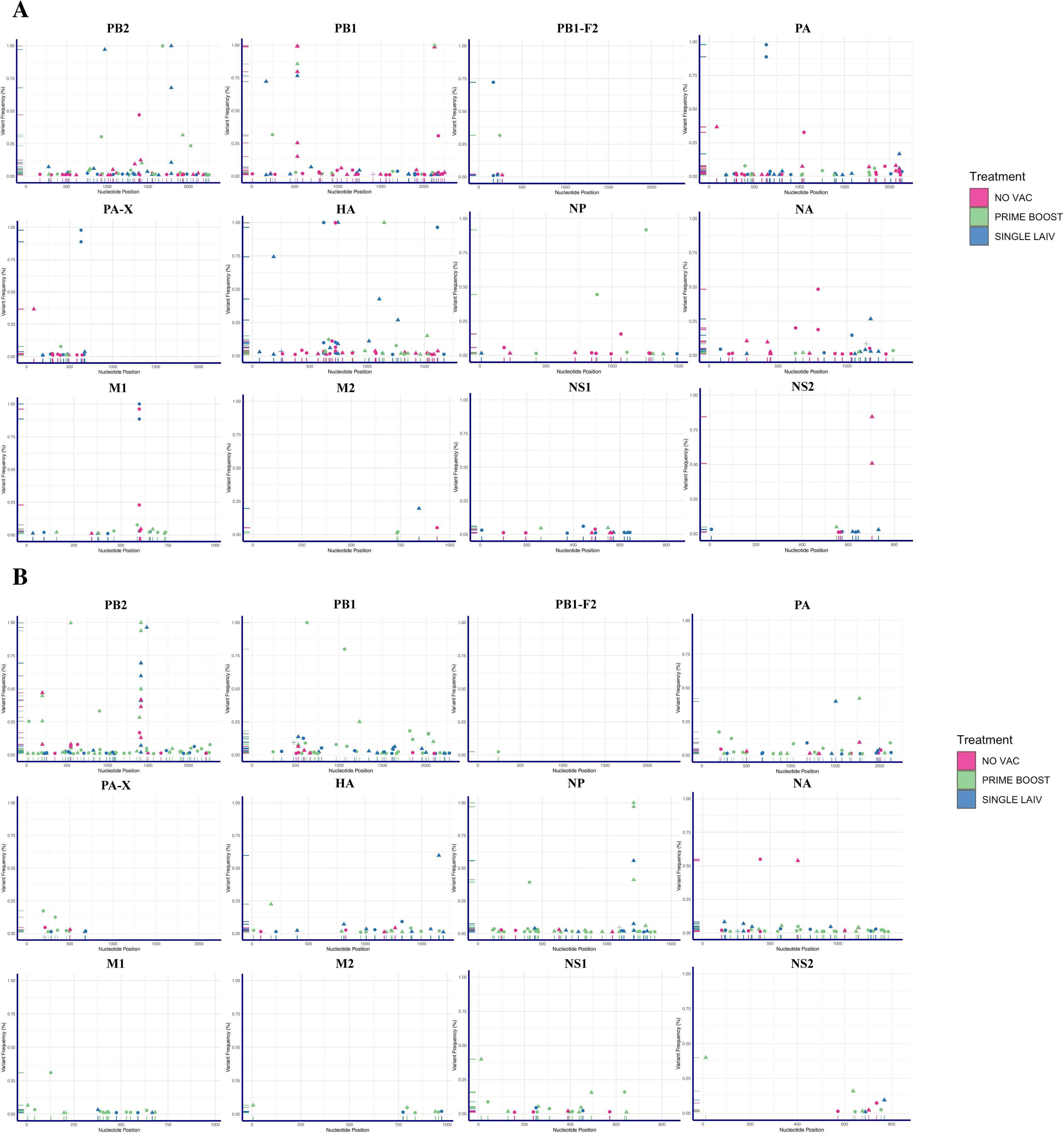
Single nucleotide variants (SNVs) of H1N1 and H3N2 viruses identified within pigs from different vaccination statuses. Summary of identified nucleotide polymorphisms of H1N1 (A) and H3N2 (B) virus in BALF samples from PRIME BOOST (light green), SINGLE LAIV (blue), and NO VAC (pink) pigs. Each unique single nucleotide variant (SNV) was annotated in either synonymous (triangle), nonsynonymous (circle) or stop-gained (cross) mutation. The x-axis represents the nucleotide site where the SNV was placed and the y-axis represents the frequency of the SNV within the virus population.

**Figure 5 - figure supplement 1.**
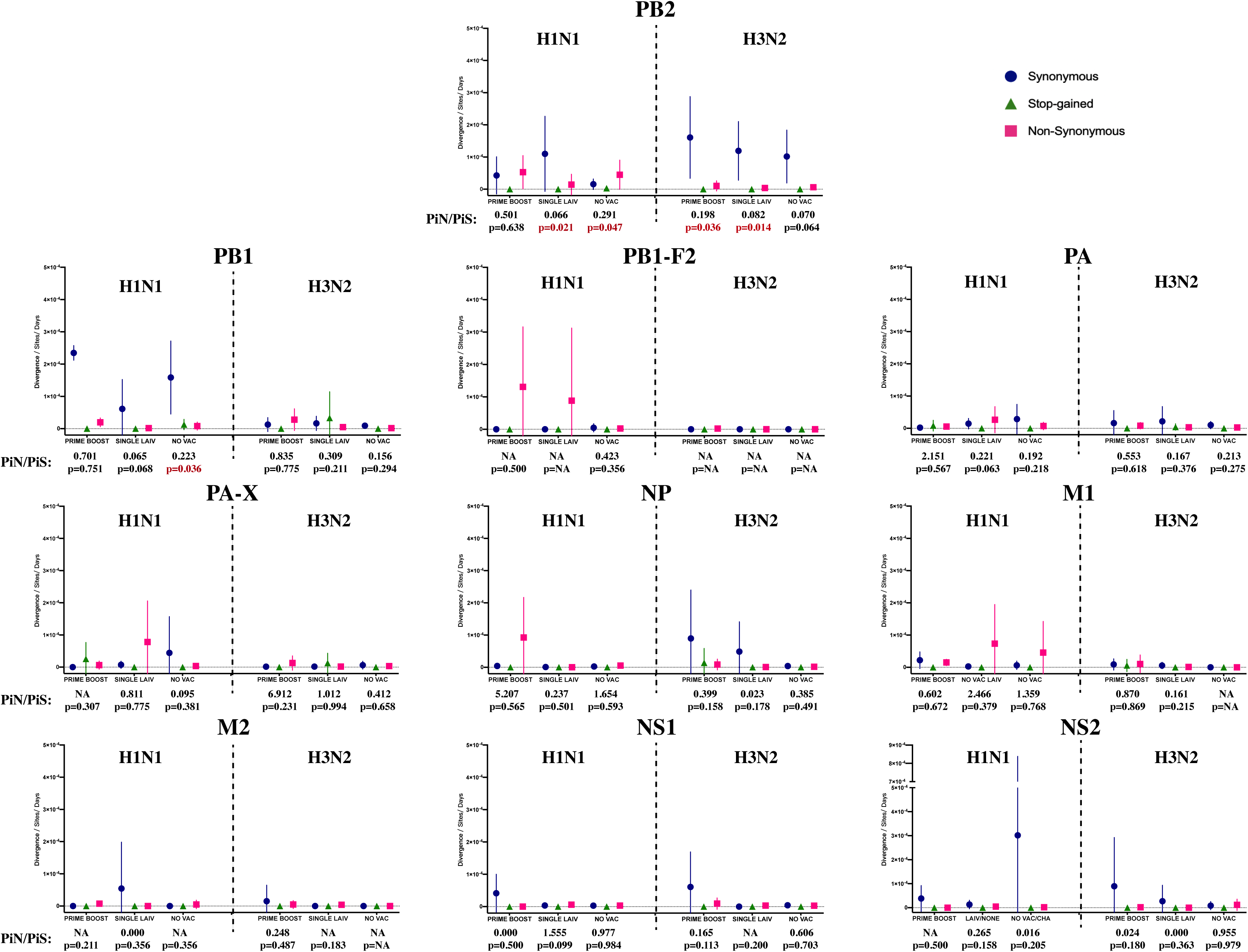
The evolutionary rate and the ratio of nonsynonymous to synonymous nucleotide diversity (PiN/PiS) of H1N1 and H3N2 influenza A virus (IAV) on coding regions located in internal genes. Figure settings and statistical analysis are the same as in Figure 5B.

**Figure 6 - figure supplement 1.**
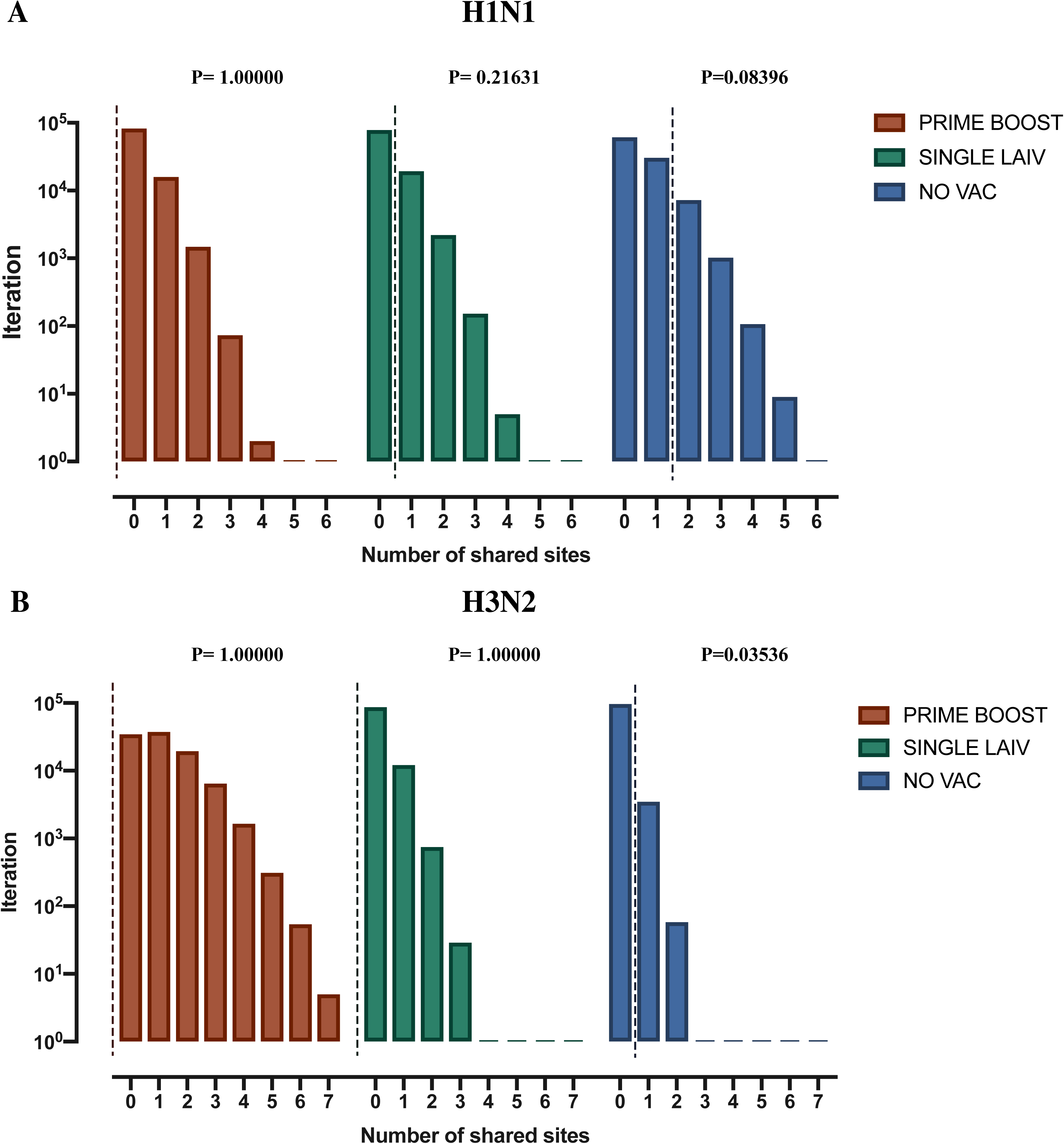
Permutation test on shared amino acid polymorphic sites in the full genome of H1N1 and H3N2 influenza A viruses (IAV). The permutation test of shared amino acid polymorphisms in H1N1 (A) and H3N2 (B) IAV genome in pigs by treatment groups. Permutation test and figure setting parameters are the same as in Figures 6B and 6D.

## Source data files

**Figure 2 – source data 1. Phylogenetic analysis of influenza plaques isolated from pigs with various vaccination statuses.** Phylogenetic trees of eight IAV genes from viral plaque isolates were constructed using the maximum likelihood method with 1000 bootstraps and best fitted nucleotide substitution model. The color of leaf nodes indicates the genetic origin of the IAV segments. The red and green taxa represent the sequences are derived from the H1N1 and H3N2 challenge viruses, respectively. The blue taxa are sequences that originated from the mixed genotype plaques. The genome of H1N1 and H3N2 challenge viruses are displayed as pink taxa and the other reference sequences are shown as black taxa. The bootstrap values are proportionally displayed by the circle size of the tree nodes.

**Figure 2 – source data 2. Influenza consensus nucleotide sequences from isolated plaques.**

**Figure 3 – source data 1. Infection dynamics of H1N1 and H3N2 challenge viruses assessed by subtype specific RRT-PCR in nasal swabs and broncho-alveolar fluid samples (BALF).**

**Figure 4 – source data 1. Summarized information of H1N1 single nucleotide variants (SNVs) identified in vaccinated and unvaccinated pigs.**

**Figure 4 – source data 2. Summarized information of H3N2 single nucleotide variants (SNVs) identified in vaccinated and unvaccinated pigs.**

## Supplementary Files

**Supplementary file 1. Number of broncho-alveolar lavage fluid (BALF) samples and number of plaques available for the study.**

**Supplementary file 2. Values of nucleotide diversity in H1N1 and H3N2 challenge viruses by coding regions and treatment groups.**

**Supplementary file 3. Spearman correlation between humoral and cellular immune responses and IAV HA nucleotide diversity in individual pigs.**

**Supplementary file 4. Detection of amino acid changes at functional sites.** The percentage of annotated functional associated single nucleotide variants (SNVs) from the total identified SNVs in H1N1 (A) and H3N2 (B) virus. The number of identified SNVs and the total number of consensus nucleotides where these SNVs were called by treatment groups are shown above the bars. Only the nonsynonymous and stop-gained mutations in our data were queried against the database of Sequence Feature Variant Types tool in the Influenza Research Database. Phenotypic summary of annotated functional related mutations in H1N1 (C) and H3N2 (D) virus. The quantity of annotated functional related mutations detected in pigs by treatment group is shown above each bar, and the mutations are colored based on their putative phenotypic effect on the virus.

**Supplementary file 5. All identified functional single nucleotide variants (SNVs) in the H1N1 virus with annotations.**

**Supplementary file 6. All identified functional single nucleotide variants (SNVs) in the H3N2 virus with annotations.**

**Supplementary file 7. Nucleotide and amino acid homology between A/swine/Minnesota/PAH-618/2011 (H1N1) and A/swine/Minnesota/080470/2015 (H3N2) viruses.**

